# Average local nucleosome motion remains nearly constant during interphase in living human cells

**DOI:** 10.64898/2026.04.29.721002

**Authors:** Yu Nagata, Shiori Iida, Masa A. Shimazoe, Sachiko Tamura, Kako Nakazato, Kai Shimizu, Yuki Hatoyama, Masato T. Kanemaki, Kazuhiro Maeshima

## Abstract

Dynamic chromatin behavior, which is related to chromatin accessibility, plays a critical role in various genome functions such as RNA transcription and DNA replication/repair. Previous studies using highly synchronized cells showed that average local chromatin motion, captured by single-nucleosome imaging on a subsecond time scale, remained nearly constant throughout G1, S, and G2 phases in living human cells. Here, we combined single-nucleosome imaging with Fucci cell-cycle probes to test this finding in asynchronous living human cells. Using HeLa and HCT116 cells expressing H2B-HaloTag and Fucci probes, we found that local nucleosome motion remained similar on average throughout interphase. Consistently, H3.3-Halo-labeled nucleosomes, which are enriched in Hi-C A compartments (euchromatin), also showed near-constant motion throughout interphase. Transcription inhibition increased nucleosome motion throughout interphase. Local nucleosome motion also increased following cellular perturbations, such as replication stress or DNA damage. Our findings suggest that near-constant chromatin motion supports housekeeping functions under similar physical conditions during interphase. They also suggest that cells can transiently change chromatin motion to perform ad hoc tasks in response to intra- and extracellular signals, such as DNA damage.

## Introduction

A long strand of genomic DNA is wrapped around core histones to form nucleosomes [1, 2]. Strings of nucleosomes associated with numerous other proteins and RNAs are organized as chromatin in cells [3, 4].

A polymer physics viewpoint suggests that chromatin has viscoelastic properties [5–8], which mean that the physical properties of chromatin can change depending on the time and size scales used for measurements. Chromatin appeared solid-like on minute/micrometer or longer spatiotemporal scales [7, 9]. In this situation, each chromosome is quite stably occupied in the territory [10] without excess intermingling and subsequent chromosome breaks, which contributes to maintaining genome integrity. On the other hand, chromatin was locally more flexible and liquid-like when measured on subsecond/100–300 nm scales [7, 11] (in vitro results, see [12]). This can increase chromatin accessibility for target searching and facilitate DNA transaction reactions such as RNA transcription and DNA replication/repair [13, 14].

How does local chromatin behavior change during interphase, from G1 to S and G2, when genomic DNA doubles [15] and the nucleus becomes larger [16–19]? Several lines of evidence have suggested that chromatin dynamics during interphase are strongly scale-dependent [20–26]. For instance, chromatin was reported to be highly dynamic in G1 and much more constrained in S and G2 phases when specific genomic loci in yeast and human cells were tracked on a 1 min or longer time scale [20–22]. However, a previous study using highly synchronized human cells showed that average nucleosome motion remained nearly constant on a shorter, subsecond time scale (up to ∼0.5 s) throughout interphase [18]. Furthermore, Repli-Histo labeling of four known chromatin classes from euchromatin to heterochromatin (IA, IB, II, and III), revealed that the more euchromatic (earlier-replicated) and more heterochromatic (later-replicated) regions exhibited greater and lesser nucleosome motions, respectively, and that the motion profile of each chromatin class persisted throughout interphase in synchronized human cells [14, 27].

However, the prolonged drug treatments used in these interphase studies, such as lovastatin [28], thymidine [29], RO-3306 [30], might directly or indirectly affect chromatin behavior. Indeed, in such synchronized cells, nuclear volumes in G2 are ∼2.5-fold larger than those in G1 [18]. To overcome this prolonged drug treatment problem, local nucleosome motion during interphase should be investigated under more physiological conditions without synchronization. To address this issue, Fucci probes, which visualize cell-cycle progression through G1, S, and G2, are powerful tools [31–35]. Fucci is a genetically encoded live-cell reporter that visualizes cell-cycle progression based on the phase-specific degradation of Cdt1- and Geminin-derived fluorescent probes. Cells in G1 are labeled in one color, whereas cells in S/G2/M are labeled in another color.

Here, we combined single-nucleosome imaging using H2B-HaloTag with Fucci probes in living cells. Single-nucleosome imaging allows detailed analyses of local chromatin behavior and is a powerful tool for sensitively detecting changes in chromatin motion in living cells [36–38]. We found that average local nucleosome motion remains nearly constant throughout interphase. Given that local chromatin motion can govern genomic DNA accessibility for target searching [13, 27] or recruitment of molecular machinery [21], our findings provide new insight into the physical behavior of chromatin during housekeeping functions in living cells, such as RNA transcription and DNA replication/repair.

## Results

### Establishment of HeLa Fucci2 cells expressing H2B-HaloTag for cell-cycle analysis

To perform single-nucleosome imaging while monitoring cell-cycle progression, we established HeLa cells expressing H2B-HaloTag together with the Fucci2 probes mCherry-hCdt1(30/120) and mVenus-hGeminin(1/110)(Fig. 1A)[32, 33]. We separately acquired images of DAPI-stained DNA, mCherry-hCdt1, and mVenus-hGeminin from each formaldehyde (FA)-fixed cell (Fig. S1A). Image analysis showed that G1-phase cells with low DAPI intensity, consistent with low DNA content, had high red fluorescence from mCherry-hCdt1. As DAPI intensity increased, green fluorescence from mVenus-hGeminin also increased, indicating progression from G1 to S and G2, and validating the Fucci2 system in this established cell line (Fig. S1B). Based on the fluorescence intensities of mCherry and mVenus (Fig. 1B), k-means clustering classified cells into four cell-cycle populations: early G1, G1, G1-S, and S-G2 (Fig. 1C). Nuclear volumes of FA-fixed G1, G1-S, and S-G2 cells were measured using confocal laser scanning microscopy. Image stacks were analyzed using Imaris software (Fig. 1D). We confirmed that, without cell synchronization, nuclear volume increased 1.66-fold from G1 (631.45 ± 78.63 μm^3^) to S-G2 phase (1047.93 ± 118.75 µm^3^) as genomic DNA doubled (Fig. 1E).

**Figure 1.**
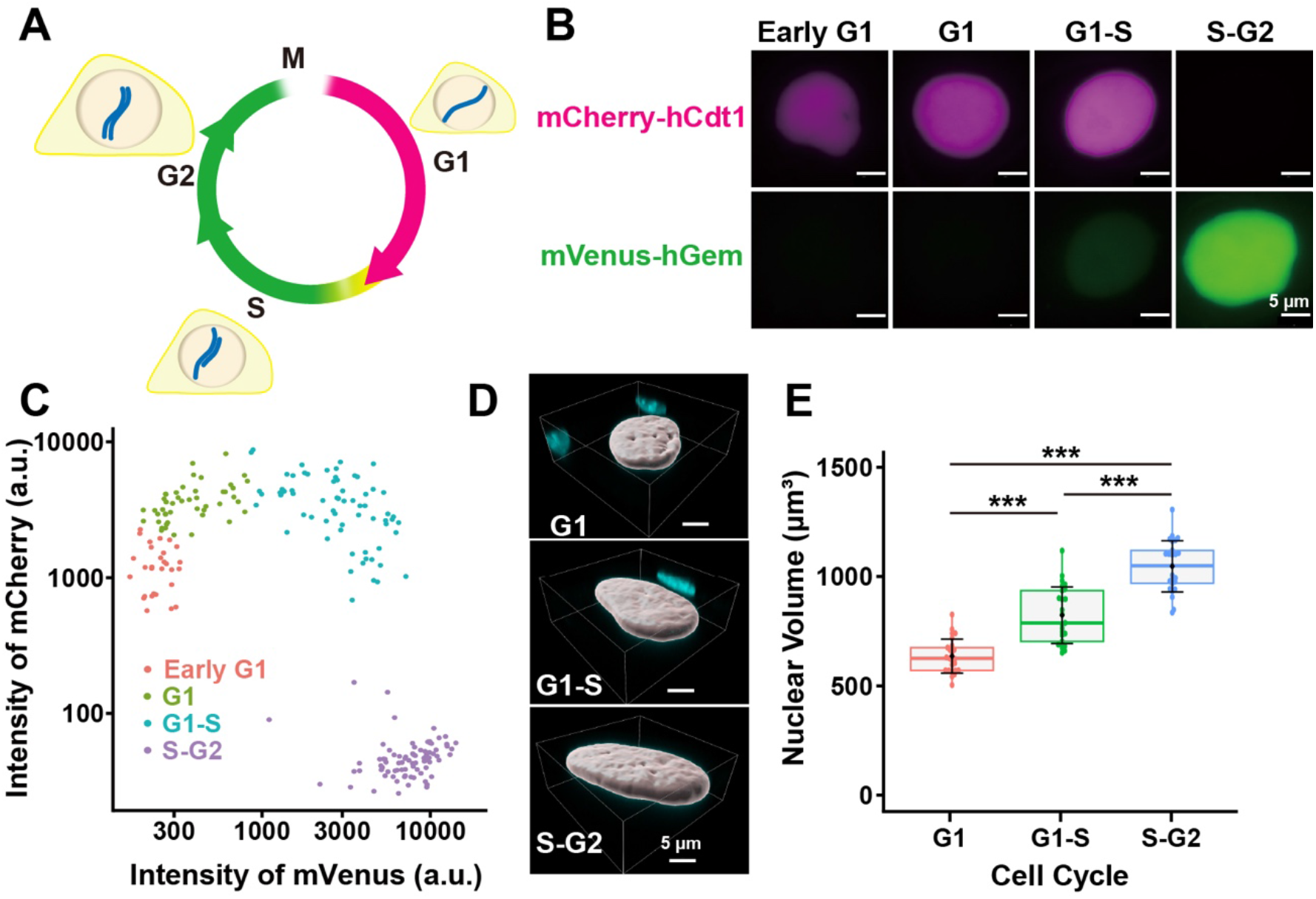
Establishment of HeLa Fucci2 cells expressing H2B-HaloTag for cell-cycle analysis. **(A)** Schematic illustration of cell-cycle progression monitored by the Fucci2 system. Cells in G1 are labeled by mCherry-hCdt1(30/120), whereas cells in S/G2/M are labeled by mVenus-hGeminin (1/110). **(B)** Representative images of Fucci2 signals in HeLa cells. mCherry-hCdt1 and mVenus-hGeminin fluorescence in early G1, G1, G1-S, and S-G2 cells is shown. Scale bars, 5 μm. **(C)** Classification of cell-cycle populations based on Fucci2 fluorescence intensities. Scatter plot of mCherry-hCdt1 and mVenus-hGeminin fluorescence intensities showing four cell-cycle populations, early G1, G1, G1-S, and S-G2, defined by k-means clustering. a.u., arbitrary units. (Early G1: n = 29 cells, G1: n = 47 cells, G1-S: n = 68 cells, S-G2: n = 82 cells). **(D)** Representative three-dimensional surface rendering of nuclei in G1, G1-S, and S-G2 cells used for nuclear volume measurement. Scale bars, 5 μm. **(E)** Nuclear volumes in G1, G1-S, and S-G2 cells (n = 25 cells per sample). Nuclear volume increased with interphase progression. Statistical significance is indicated by asterisks (***, P < 0.0001). Mean ± SD values are: G1, 631.45 ± 78.63 μm^3^; G1-S, 820.78 ± 131.36 μm^3^; S-G2, 1047.93 ± 118.75 μm^3^. *P* values from Holm-adjusted Wilcoxon rank sum tests were 1.89 × 10^-7^ for G1 versus G1-S, 2.26 × 10^-7^ for G1-S versus S-G2, and 4.75 × 10^-14^ for S-G2 versus G1.

We established HeLa Fucci2 cells stably expressing HaloTag-labeled core histone H2B (H2B-Halo), which was visualized using the HaloTag ligand JF646 [39] (Figs. 2A and S2A). H2B-Halo appeared to be properly incorporated into nucleosomes genome-wide, including both euchromatic and heterochromatic regions (Figs. 2A and S2B), likely because histone H2B turns over within a few hours [40].

**Figure 2.**
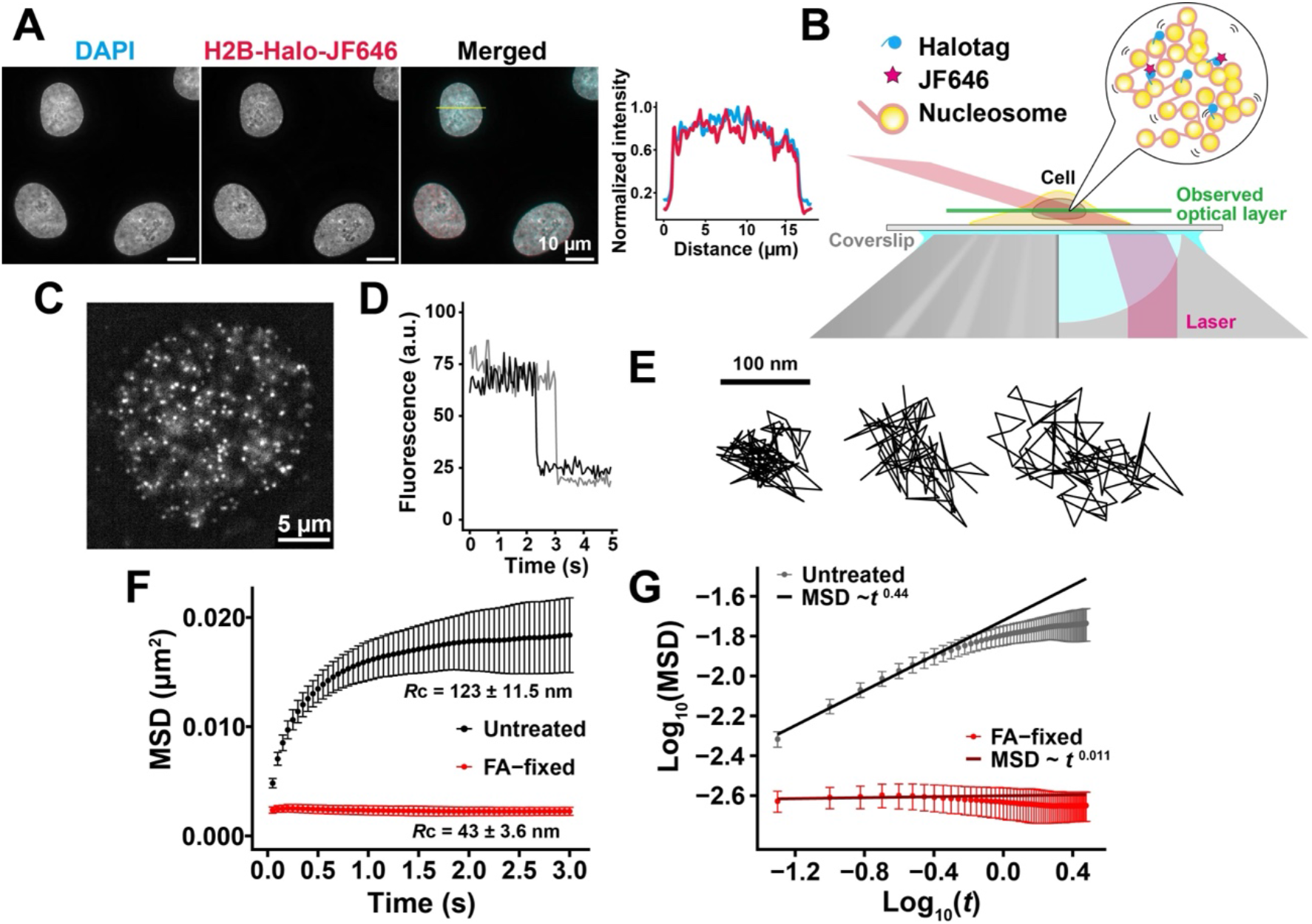
Single-nucleosome imaging of H2B-HaloTag–JF646 in living HeLa cells. **(A)** Representative images of DAPI-stained DNA, H2B-HaloTag labeled with JF646, and the merged image in a HeLa cell. Scale bars, 10 μm. **(B)** Schematic illustration of single-nucleosome imaging by oblique illumination microscopy (HILO). H2B-HaloTag in nucleosomes was sparsely labeled with JF646 and imaged within a thin optical layer in the nucleus by oblique illumination or HILO [41, 42]. **(C)** Representative image of single H2B-HaloTag–JF646 dots in the nucleus of a living HeLa cell after background subtraction. Scale bar, 5 μm. **(D)** Representative fluorescence intensity traces of single H2B-HaloTag–JF646 dots showing single-step photobleaching. a.u., arbitrary units. **(E)** Three representative trajectories of single nucleosomes in a living HeLa cell. Scale bar, 100 nm. **(F)** Mean square displacement (MSD) plots of H2B-HaloTag–JF646 dots in untreated living cells (Nucleosome trajectories used per cell: 507 ∼ 1363; n = 31 cells) and formaldehyde (FA)-fixed cells (Nucleosome trajectories used per cell: 243 ∼ 985; n = 31 cells). Nucleosome motion was strongly suppressed in FA-fixed cells. Error bars represent SD among cells. **(G)** Log-log plot of MSD versus time interval in untreated living cells and FA-fixed cells. The slope was approximately 0.44 in untreated cells and 0.011 in FA-fixed cells, indicating subdiffusive nucleosome motion in living cells and nearly immobilized nucleosomes in FA-fixed cells.

We then sparsely labeled H2B-Halo [18] with JF646 [39] to visualize single nucleosomes and performed live-cell imaging using oblique illumination microscopy (HILO), which illuminates a thin layer within the nucleus (Fig. 2B)[41, 42]. JF646-labeled nucleosomes were recorded at 50 ms/frame in asynchronous HeLa cells (∼100 frames, 5 s total) and observed as clear dots (Fig. 2C; Movie S1). These dots showed a single-step photobleaching profile (Fig. 2D), suggesting that each dot represents a single nucleosome labeled with H2B-Halo-JF646. The individual dots were fitted with a 2D Gaussian function to estimate the precise positions of nucleosomes [43, 44]. They were tracked using u-track software [45](Movie S2) to obtain their motion trajectories (Fig. 2E). The position determination accuracy was 9.33 nm (Fig. S2C). From the obtained nucleosome trajectory data (e.g., Fig. 2E), we calculated mean square displacement (MSD) (Figs. 2F and S2D), which shows how nucleosomes move over a certain time period. Tracking was specific to nucleosome-incorporated H2B-Halo-JF646, as the small free H2B-Halo pool (4.4%) diffused too fast to be observed (Fig. S2E).

MSD plots appeared sub-diffusive (Fig. 2F). Chemical fixation (FA) of the cells almost immobilized the JF646-labeled nucleosomes. MSD neared a plateau at ∼ 3 s (Fig. 2F), which is proportional to the square of the radius of constraint [Rc; P (plateau value) = 6/5 × Rc^2^; [46]]. The estimated radius of the nucleosome motion constraint was 123 ± 11.5 nm, which is similar to the chromatin domain size with a diameter of 100–200 nm [47–51] and is also consistent with our previous studies [18, 52]. A log-log plot of the MSD data showed that the MSD of local nucleosome motion for the first 0.5 sec was proportional to t^0.44^ (Fig. 2G). Beyond 0.5 s, the plot could not be fitted linearly in the double logarithmic graph (Fig. 2G), indicating that the motion mode differs past 0.5 s. We therefore focused on the short time range up to 0.5 s in the following analyses, as in our previous studies [18, 51–53].

### Average local nucleosome motion remains nearly constant throughout interphase

Next, we examined local nucleosome motion during interphase. We performed single-nucleosome imaging in living cells and then obtained mCherry-hCdt1 and mVenus-hGeminin images from the same cells. Based on the fluorescence intensities of mCherry and mVenus, k-means clustering classified cells into four cell-cycle populations: early G1, G1, G1-S, and S-G2. Nucleosome motion in each phase was quantified by MSD. MSD plots for the G1, G1-S, and S-G2 phases were similar, with no significant differences (Fig. 3A), consistent with our previous studies [18]. In contrast, the early G1 phase showed slightly higher MSD values than the other phases (Fig. 3A). A scatter plot of MSD versus normalized mVenus fluorescence intensity showed that MSD was slightly higher in the low-mVenus region corresponding to early G1 (Fig. S3A), whereas it remained nearly constant over the rest of interphase (Fig. 3B). Taken together, these results indicate that average local nucleosome motion remains nearly constant throughout interphase.

**Figure 3.**
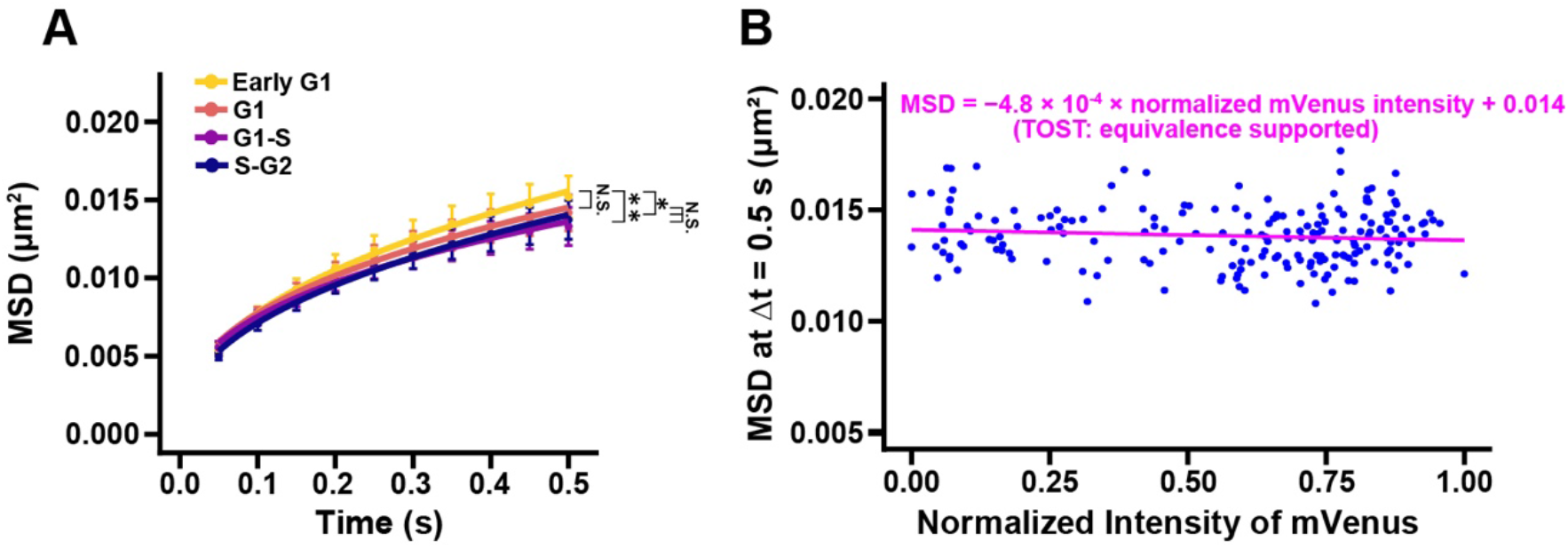
Average local nucleosome motion remains nearly constant throughout interphase. **(A)** Mean square displacement (MSD) plots of single nucleosomes labeled by H2B-Halo in early G1 (n = 25 cells), G1 (n = 30 cells), G1-S (n = 30 cells), and S-G2 (n = 30 cells) cells classified by Fucci2 signals. MSD values were similar among G1, G1-S, and S-G2 cells, whereas early G1 cells showed slightly higher MSD values. Error bars represent SD among cells. Nucleosome trajectories used per cell: 766 ∼ 2369. Holm-adjusted Kolmogorov-Smirnov test *P* values were 0.153 for early G1 versus G1, 6.84 × 10⁻^4^ for early G1 versus G1-S, 1.66 × 10⁻^2^ for early G1 versus S-G2, 0.138 for G1 versus G1-S, 0.786 for G1 versus S-G2, and 0.786 for G1-S versus S-G2. N.S., not significant. (*, *P* < 0.05; **, *P* < 0.001) **(B)** Scatter plot of MSD at Δt = 0.5 s versus normalized mVenus fluorescence intensity. When early G1 cells were excluded, MSD remained nearly constant over the rest of interphase, as indicated by the small regression slope and two one-sided tests (TOST) analysis. n = 197 cells.

So far, we focused on average nucleosome motion revealed by H2B-Halo-JF646 imaging in the early G1, G1, G1-S, and S-G2 populations. It is possible that heterogeneity in nucleosome motion was masked by population averaging. To examine this possibility, we analyzed the MSD distributions at Δt = 0.5 s using Richardson-Lucy (RL) deconvolution analysis, which resolves heterogeneous MSD distributions into multiple motional components [11, 54]. RL analysis identified four components, designated super-slow, slow, fast, and super-fast motions, in all cell-cycle phases (Fig. S3B). The overall MSD distribution profiles were similar among G1, G1-S, and S-G2, whereas early G1 showed a modest shift of “slow” and “fast” components toward higher mobility (Fig. S3B). Thus, both MSD and RL analyses support the conclusion that average local nucleosome motion and the overall MSD distribution remain nearly constant throughout interphase.

To further examine nucleosome motion in a more defined chromatin population, we next examined H3.3-Halo nucleosomes, which are enriched in active/euchromatic regions [55–57]. We stably expressed H3.3-Halo in HeLa Fucci2 cells (Fig. S4A) [51]. Expressed H3.3-Halo was localized to DAPI-poor regions in nuclei (Fig. 4A). Pull-down of H3.3-Halo-incorporated nucleosomes and genomic sequencing confirmed enrichment in Hi-C A compartments with active histone marks, and depletion from repressive marks (Fig. S4B)[51]. Single-nucleosome imaging of H3.3-Halo nucleosomes (Figs. 4B and S4C-E) in HeLa Fucci2 cells (Fig. 4C) showed that H3.3-Halo nucleosomes were more mobile than H2B-Halo nucleosomes during interphase (Fig. 4D; Movie S3). Average H3.3-Halo nucleosome motion remained nearly constant throughout interphase (Figs. 4E and S4F). These findings strengthen the conclusion that local nucleosome motion remains nearly constant throughout interphase.

**Figure 4.**
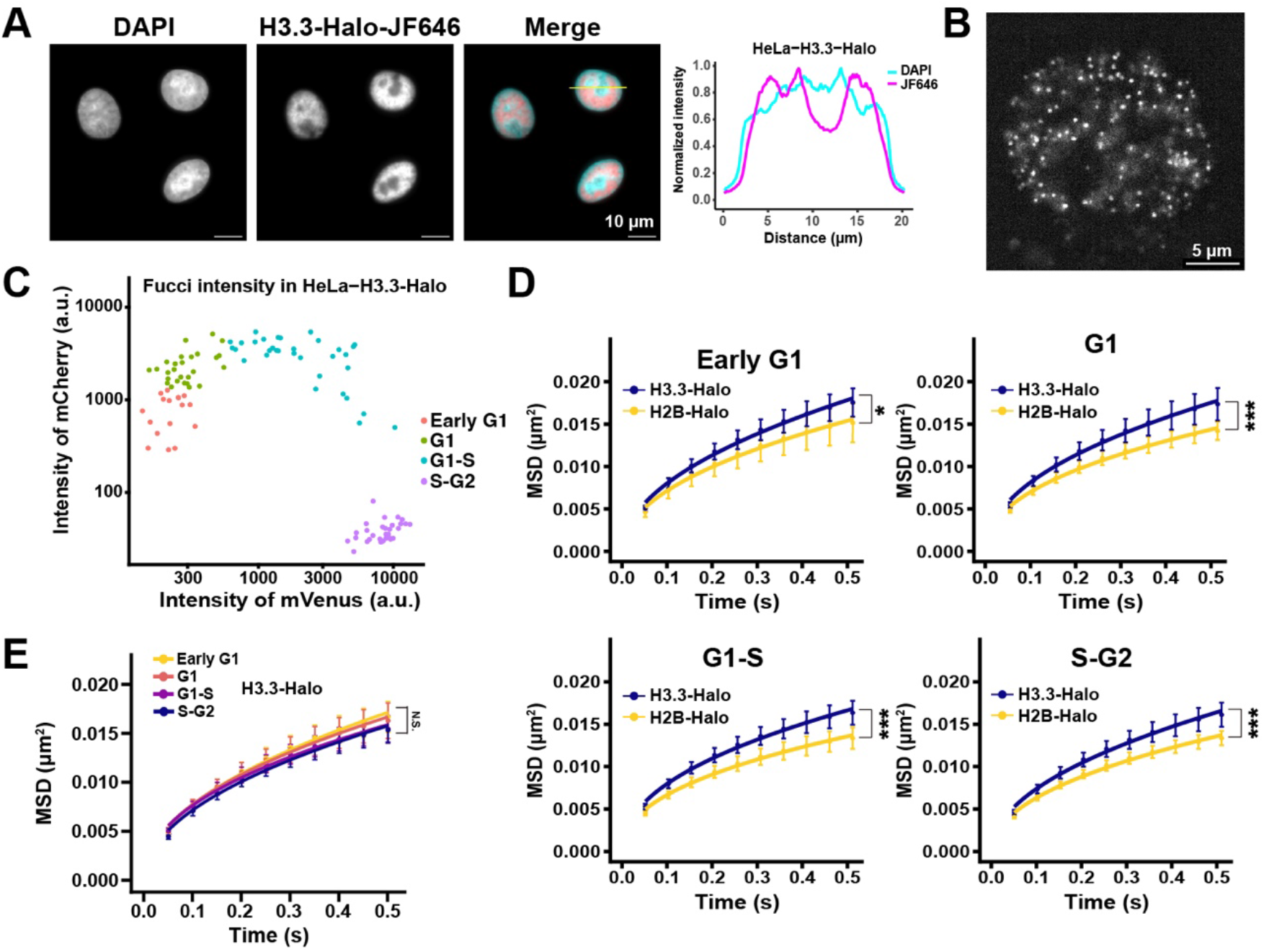
H3.3-Halo nucleosomes show greater but nearly constant motion throughout interphase. **(A)** Representative images of DAPI-stained DNA, H3.3-Halo-JF646, and merged images in HeLa Fucci2 cells. Normalized fluorescence intensity profiles of DAPI and H3.3-Halo-JF646 along the yellow line are shown on the right. Scale bars, 10 μm. **(B)** Representative image of single H3.3-Halo-JF646 dots in the nucleus of a living HeLa Fucci2 cell after background subtraction. Scale bar, 5 μm. **(C)** Classification of H3.3-Halo-expressing HeLa cells into early G1, G1, G1-S, and S-G2 populations based on Fucci2 fluorescence intensities. Early G1: n = 17 cells; G1: n = 27 cells; G1-S: n = 33 cells; S-G2: n = 31 cells. a.u., arbitrary units. **(D)** MSD plots of H3.3-Halo and H2B-Halo nucleosomes in early G1 (n = 17 cells), G1 (n = 20 cells), G1-S (n = 20 cells), and S-G2 (n = 20 cells) cells. H3.3-Halo nucleosomes showed significantly greater motion than H2B-Halo nucleosomes in all cell-cycle populations examined. Kolmogorov-Smirnov test *P* values were 0.0156 for early G1, 1.33 × 10⁻⁶ for G1, 1.33 × 10⁻⁶ for G1-S, and 1.43 × 10⁻⁷ for S-G2 (*, *P* < 0.05; ***, *P* < 0.0001). H3.3-Halo nucleosome trajectories used per cell: 569–2300. H2B-Halo nucleosome trajectories used per cell: 655 ∼ 1957. **(E)** MSD plots of H3.3-Halo nucleosomes in early G1, G1, G1-S, and S-G2 cells. MSD values were similar among the four cell-cycle populations, with no significant differences. Holm-adjusted Kolmogorov-Smirnov test *P* values were 1.00 for early G1 versus G1, 0.250 for early G1 versus G1-S, 0.0663 for early G1 versus S-G2, 1.00 for G1 versus G1-S, 0.168 for G1 versus S-G2, and 1.00 for G1-S versus S-G2. N.S., not significant. Data in (D) and (E) represent mean ± SD among cells.

### Transcription inhibition increases average nucleosome motion throughout interphase of HeLa cells

It has been reported that transcription inhibition by various inhibitors (e.g., α-amanitin, DRB, triptolide, and THZ1) or rapid RNA Pol II depletion increases nucleosome motion in living cells [48, 52, 58–64], suggesting that transcription globally constrains chromatin. While average nucleosome motion remains nearly constant throughout interphase, as demonstrated above, we wondered whether transcription constrains chromatin throughout interphase or only in specific phases. To address this question, we examined how transcription inhibition affects local nucleosome motion during interphase. To inhibit transcription, cells were treated with THZ1 [65], a covalent CDK7 inhibitor that suppresses phosphorylation (Ser5) of RNA polymerase II (Fig. 5A) and subsequent transcription. Nucleosome motion was quantified by MSD in early G1, G1, G1-S, and S-G2 phases (Fig. S5A). Transcription inhibition by THZ1 increased MSD values in all interphase phases examined, although the effect was less prominent in G1-S than in the other phases (Fig. 5B). Thus, transcription inhibition increases average nucleosome motion not only in a specific phase but throughout interphase, suggesting that RNA Pol II-dependent transcriptional activity contributes to constraining local nucleosome motion in living cells.

**Figure 5.**
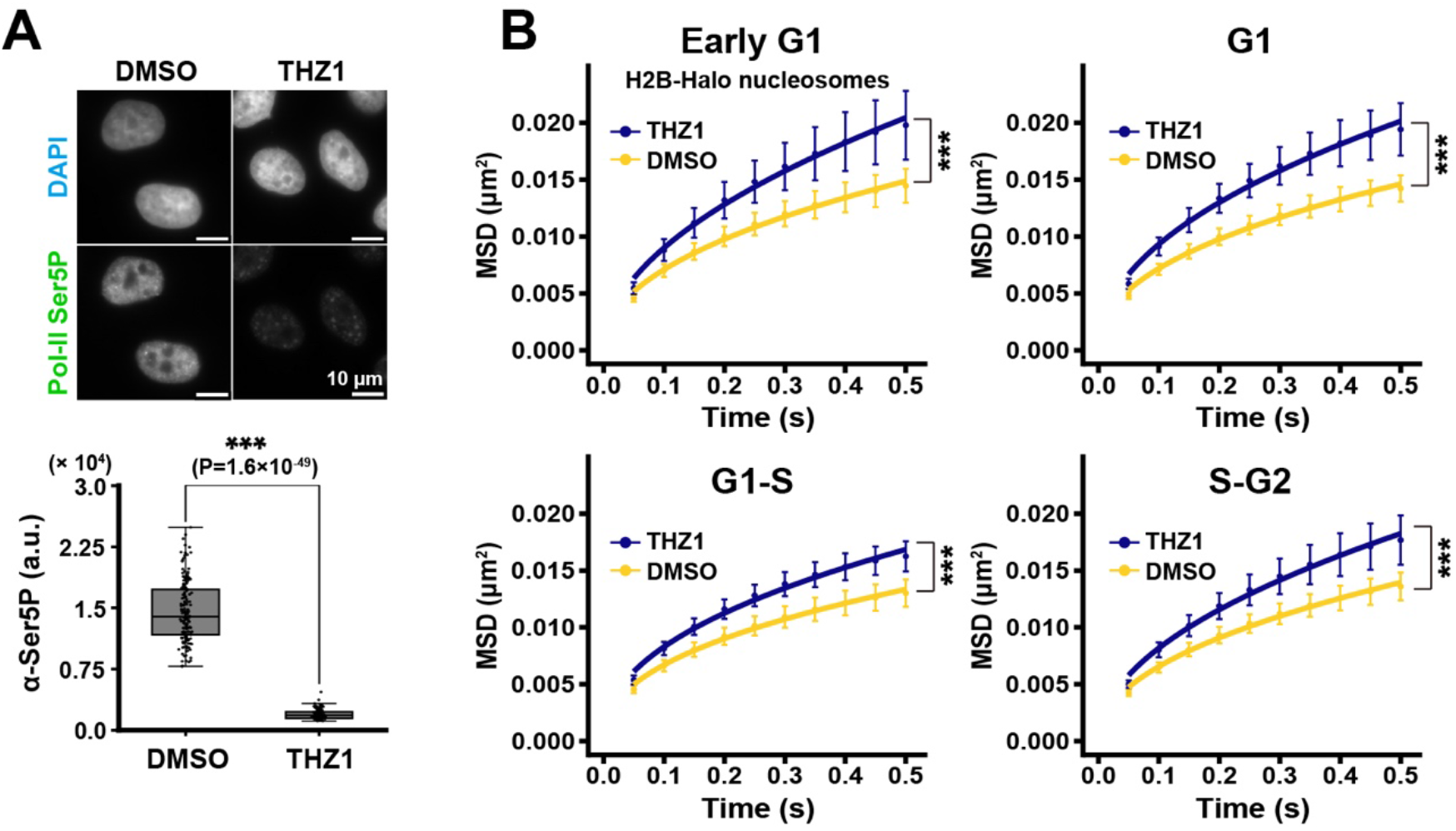
Transcription inhibition increases average nucleosome motion throughout interphase. **(A)** Validation of transcription inhibition by THZ1. Representative images of DAPI and RNA polymerase II Ser5 phosphorylation (Pol II Ser5P) in DMSO- and THZ1-treated HeLa cells are shown together with quantification of the integrated Pol II Ser5P intensity. DMSO: n = 166 cells; THZ1: n = 131 cells. 1 µM THZ1 treatment for 2 h markedly reduced Pol II Ser5P signals (Wilcoxon rank sum test, *P* = 1.60 × 10^-49^; ***, *P* < 0.0001). a.u., arbitrary units. Scale bar, 10 μm. **(B)** Mean square displacement (MSD) plots of single nucleosomes labeled by H2B-Halo in early G1 (DMSO: n = 25, THZ1: n = 15), G1 (DMSO: n = 24, THZ1: n = 30), G1-S (DMSO: n = 30, THZ1: n = 30), and S-G2 (DMSO: n = 30, THZ1: n = 30) cells treated with DMSO or THZ1. THZ1 significantly increased MSD in early G1, G1, G1-S, and S-G2 cells (Kolmogorov-Smirnov test *P* values were 2.39 × 10^-8^, 2.04 × 10^-12^, 5.79 × 10^-13^, and 9.24 × 10^-11^, respectively; ***, *P* < 0.0001). Error bars represent SD among cells. Nucleosome trajectories used per cell: 712 ∼ 2628.

### Replication stress and DNA damage increase average nucleosome motion

Because a previous study showed that thymidine block increased average nucleosome motion [18], we wondered whether cellular perturbations such as replication stress and DNA damage also increase average nucleosome motion. To address this issue, we treated cells with 1 mM hydroxyurea (HU) for 24 h. HU is an inhibitor of ribonucleotide reductase and is widely used to induce replication stress and S-phase arrest [66]. As in thymidine treatment, HU-treated cells accumulated around S phase (S-G2) (Fig. S5B), and their MSD increased significantly (Fig. 6A).

**Figure 6.**
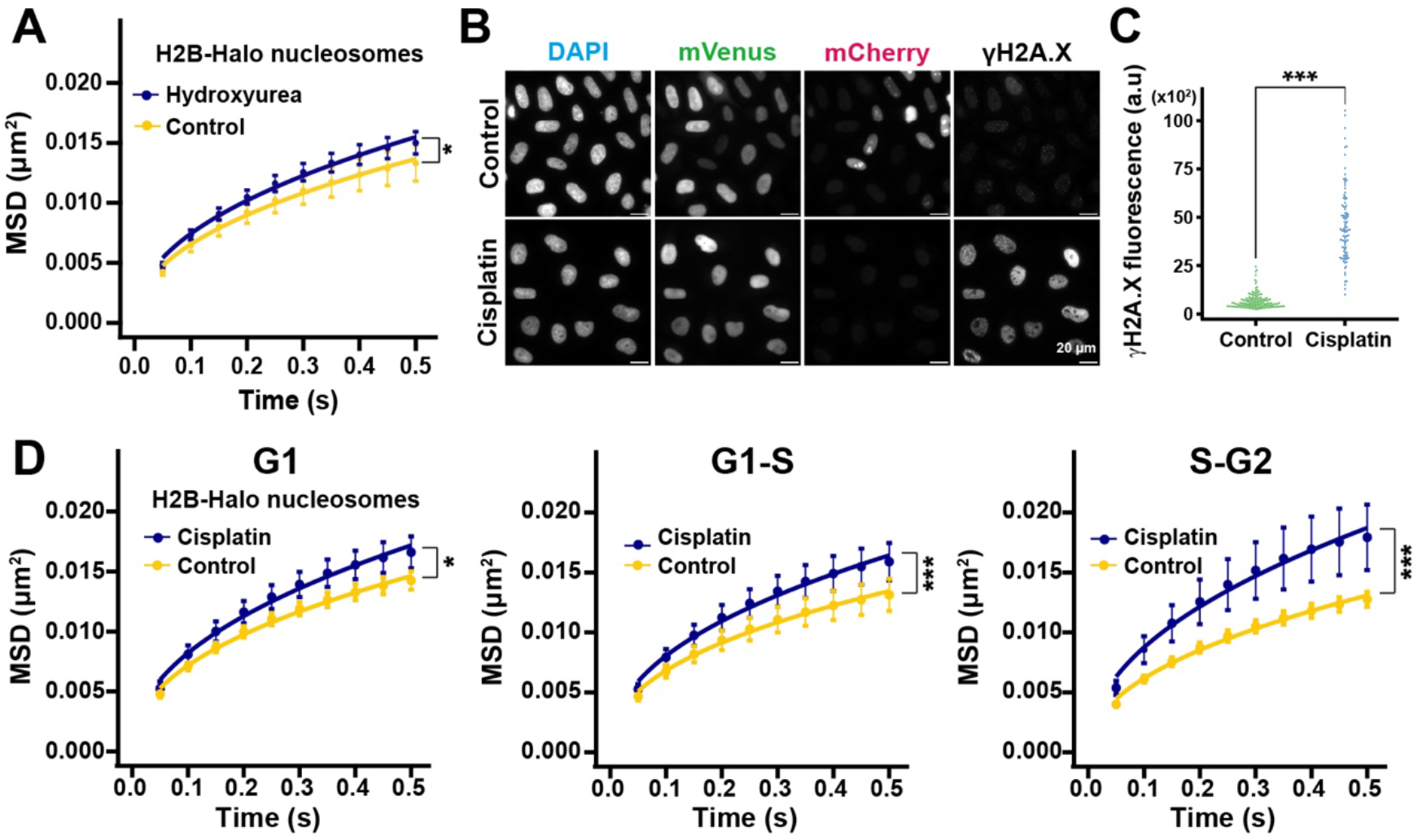
Replication stress and DNA damage increase average nucleosome motion. **(A)** Mean square displacement (MSD) plots of single nucleosomes labeled by H2B-Halo in 1mM hydroxyurea-treated (24 h) and control HeLa cells. Hydroxyurea increased nucleosome motion (Kolmogorov-Smirnov test *P* value was 3.97 × 10^-3^, *, *P* < 0.05). Error bars represent SD among cells. Nucleosome trajectories used per cell: 1090 ∼ 2973; n = 20 cells per sample. **(B)** Representative images of DAPI, mVenus, mCherry, and γH2A.X in control and 4 µM cisplatin-treated (24 h) HeLa cells. Scale bars, 20 μm. **(C)** Quantification of γH2A.X fluorescence intensity in control and cisplatin-treated cells. Cisplatin treatment markedly increased γH2A.X signals (Wilcoxon rank sum test, P = 2.20 × 10^-16^; ***, P < 0.0001). a.u., arbitrary units. Control: n = 145 cells, Cisplatin: n = 96 cells. **(D)** MSD plots of single nucleosomes labeled by H2B-Halo in G1 (Control: n = 20, Cisplatin: n = 3), G1-S (Control: n = 20, Cisplatin: n = 10), and S-G2 (Control: n = 20, Cisplatin: n = 20) cells treated with cisplatin or control medium. Cisplatin increased nucleosome motion in all phases examined, with the largest increase in S-G2 cells. Error bars represent SD among cells. Kolmogorov-Smirnov test *P* values were 4.52 × 10^-3^ for G1, 3.04 × 10^-5^ for G1-S, and 5.80 × 10^-10^ for S-G2 (*, *P* < 0.05; ***, *P* < 0.0001). Nucleosome trajectories used per cell: 968 ∼ 4366.

We also treated cells with 4 µM cisplatin for 24 h. Cisplatin is a DNA-damaging agent that forms intra- and interstrand DNA adducts/crosslinks, thereby interfering with DNA replication and repair [67]. We found that cisplatin-treated cells accumulated in S-G2 (Fig. S5C), showed increased γH2A.X foci, a DNA damage marker (Fig. 6B-C)[68], and exhibited a significant increase in MSD (Fig. 6D). These findings indicate that cellular perturbations including replication stress and DNA damage increase average nucleosome motion.

### Average local nucleosome motion remains nearly constant in human HCT116 cells

To test whether the properties observed above were specific to HeLa cells, we used human HCT116 cells carrying the Fucci(SA)5 system [34] (Fig. S6A-B). The HeLa cells used in this study had an abnormal karyotype (69 ± 4.2 chromosomes; n = 39 cells) [18], whereas HCT116 cells are human colon carcinoma cells with a near-diploid karyotype of 45 chromosomes [69] and have been widely used in recent genomics studies, including 4D Nucleome analyses.

We first stably expressed H2B-Halo in HCT116 Fucci(SA)5 cells (Figs. 7A and S6C-D). Their nuclear volume increased 1.49-fold from G1 (617.56 ± 144.15 µm^3^) to S-G2 phase (918.43 ± 140.02 µm^3^) as genomic DNA doubled (Fig. 7B-E). We then performed single-nucleosome imaging in living HCT116 cells (Figs. 8A-B; S7A; Movies S4-S5). The position determination accuracy was 10.43 nm (Fig. S7B). We obtained AzaleaB5-hCdt1 and h2-3-hGeminin images from the same cells to classify them into early G1, G1, G1-S, and S-G2 phases. MSD plots for these phases were similar, with no significant differences (Fig. 8C). The scatter plot of MSD versus normalized h2-3 fluorescence intensity showed only a slight increase in early G1 (Fig. S7C), whereas it remained nearly constant over the rest of interphase (Fig. 8D). Taken together, these observations in HCT116 cells support the conclusion that average local nucleosome motion remains nearly constant throughout interphase.

**Figure 7.**
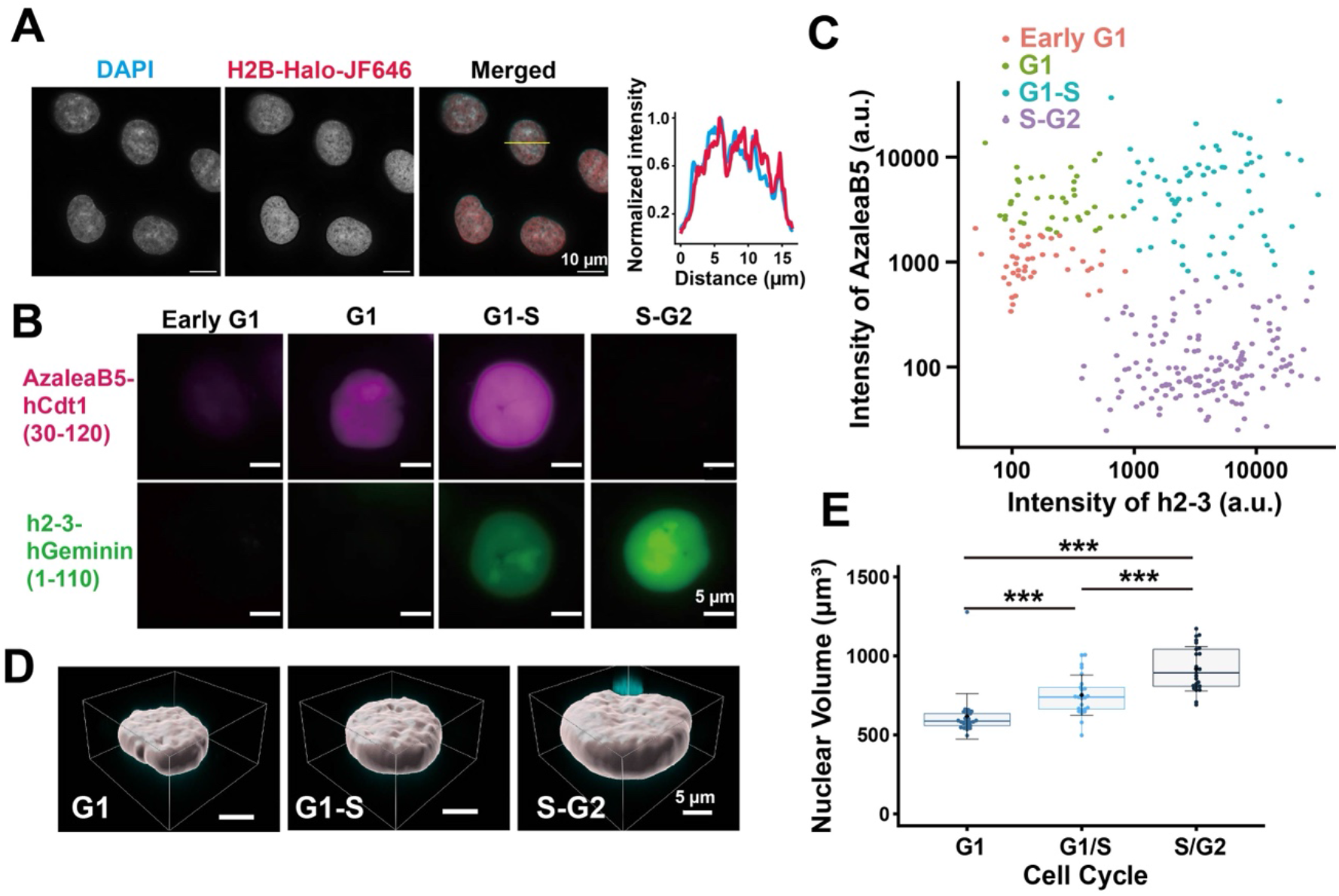
Establishment of HCT116 Fucci(SA)5 cells expressing H2B-HaloTag for cell-cycle analysis. **(A)** Representative images of DAPI-stained DNA, H2B-HaloTag labeled with JF646, and the merged image in HCT116 cells. A representative line-scan profile of DAPI and JF646 signals is shown on the right. Scale bars, 10 μm. **(B)** Representative Fucci(SA)5 fluorescence images of HCT116 cells in early G1, G1, G1-S, and S-G2 phases. AzaleaB5-hCdt1(30/120) and h2-3-hGeminin(1/110) signals are shown. Scale bars, 5 μm. **(C)** Classification of cell-cycle populations based on Fucci(SA)5 fluorescence intensities. Scatter plot of AzaleaB5-hCdt1 and h2-3-hGeminin fluorescence intensities showing four cell-cycle populations, early G1, G1, G1-S, and S-G2. a.u., arbitrary units. Early G1: n = 43 cells, G1: n = 42 cells, G1-S: n = 71 cells, S-G2: n = 146 cells. **(D)** Representative three-dimensional surface rendering of nuclei in G1, G1-S, and S-G2 cells used for nuclear volume measurement. Scale bars, 5 μm. **(E)** Nuclear volumes in G1, G1-S, and S-G2 cells. (n = 25 cells per sample) Nuclear volume increased with interphase progression. Mean ± SD values are: G1, 617.56 ± 144.15 μm^3^; G1-S, 751.52 ± 127.21 μm^3^; S-G2, 918.43 ± 140.02 μm^3^. *P* values were 1.35 × 10^-6^ for G1 versus G1-S, 3.39 × 10^-5^ for G1-S versus S-G2, and 4.41 × 10^-10^ for S-G2 versus G1. ***, *P* < 0.0001 from Holm-adjusted Wilcoxon rank sum test.

**Figure 8.**
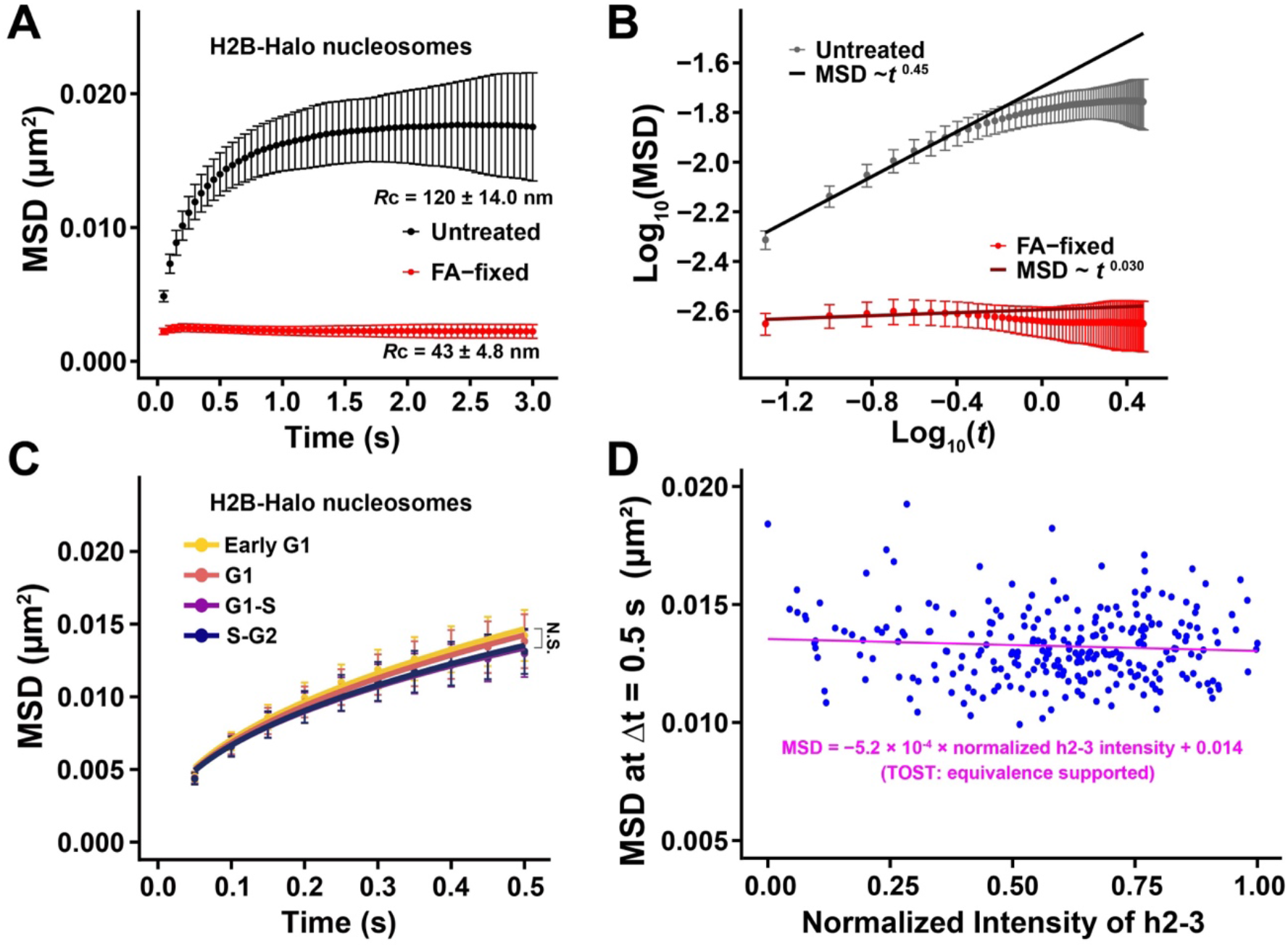
Average local nucleosome motion remains nearly constant in HCT116 cells. **(A)** Mean square displacement (MSD) plots of single nucleosomes labeled by H2B-Halo in untreated living HCT116 cells (Nucleosome trajectories used per cell: 272 ∼ 1049; n = 32 cells) and formaldehyde (FA)-fixed HCT116 cells (Nucleosome trajectories used per cell: 314 ∼ 825; n = 39 cells). Nucleosome motion was strongly suppressed in FA-fixed cells. Error bars represent SD among cells. **(B)** Log-log plot of MSD versus time interval in untreated living HCT116 cells and FA-fixed HCT116 cells. The fitted slope was approximately 0.45 in untreated cells and 0.030 in FA-fixed cells. **(C)** MSD plots of single nucleosomes labeled by H2B-Halo in early G1, G1, G1-S, and S-G2 HCT116 cells classified by Fucci(SA)5 signals. MSD values were similar among the phases, with no significant differences. Error bars represent SD among cells. Nucleosome trajectories used per cell: 364 ∼ 1435; n = 30 cells per sample. Holm-adjusted Kolmogorov-Smirnov test *P* values were 1.00 for early G1 versus G1, 9.39 × 10⁻^2^ for early G1 versus G1-S, 5.40 × 10⁻^1^ for early G1 versus S-G2, 0.354 for G1 versus G1-S, 1.00 for G1 versus S-G2, and 1.00 for G1-S versus S-G2. N.S., not significant. **(D)** Scatter plot of MSD at Δt = 0.5 s versus normalized h2-3 fluorescence intensity in HCT116 cells. MSD showed little change over the rest of interphase, as supported by the small regression slope and TOST analysis. n = 259 cells.

### Transcription inhibition increases average nucleosome motion in HCT116 cells

We next examined whether transcription inhibition similarly affects local nucleosome motion in HCT116 cells. Cells were treated with THZ1 for 2 h, and nucleosome motion was quantified by MSD in early G1, G1, G1-S, and S-G2 phases (Fig. S8). As observed in HeLa cells, THZ1 treatment increased MSD values in all phases compared with the DMSO control, although the effect was less prominent in G1-S and S-G2 than in the other phases (Fig. 9). These results support the conclusion that RNA Pol II-dependent transcriptional activity contributes to constraining local nucleosome motion in human cells.

**Figure 9.**
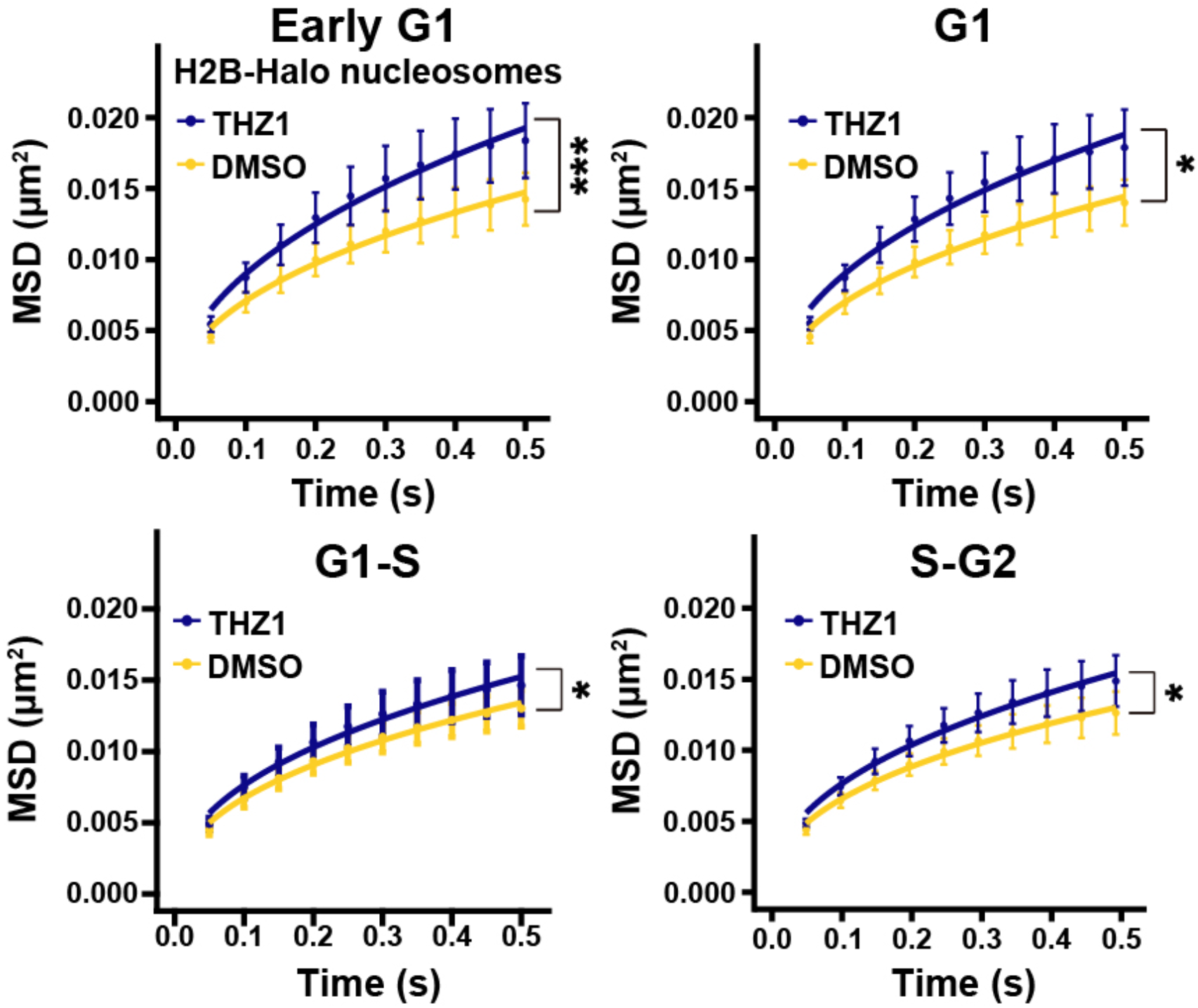
Transcription inhibition increases average nucleosome motion in HCT116 cells. Mean square displacement (MSD) plots of single nucleosomes labeled by H2B-Halo in early G1, G1, G1-S, and S-G2 HCT116 cells treated with THZ1 or DMSO. THZ1 increased nucleosome motion in all interphase phases examined. The cells were treated with 1 µM THZ1 for 2 h. Error bars represent SD among cells. Kolmogorov-Smirnov test *P* values were 9.55 × 10^-6^ for early G1, 1.16 × 10^-3^ for G1, 1.16 × 10^-2^ for G1-S, and 3.97 × 10^-3^ for S-G2; *, *P* < 0.05; ***, *P* < 0.0001. Nucleosome trajectories used per cell: 415 ∼ 1658; n = 20 cells per sample.

## Discussion

In this study, we examined local motion of H2B-Halo-labeled nucleosomes during interphase of living human cells without drug-induced cell-cycle synchronization. Our main finding is that average local nucleosome motion remains nearly constant throughout interphase (Fig. 10). Consistently, we obtained a similar result using H3.3-Halo-labeled nucleosomes, which are enriched in Hi-C A compartments with active histone marks (euchromatin) (Figs. 4E and S4F). The obtained results are also consistent with our previous study using highly synchronized HeLa and HCT116 cells [18], and indicate that the near-constant local chromatin motion observed during G1, S, and G2 is not simply an artifact caused by prolonged drug treatment for cell synchronization. In addition, Fucci intensity-based analysis allowed us to examine nucleosome motion along continuous cell-cycle progression in asynchronous cells, rather than only in discrete synchronized populations. The present results therefore strengthen the view that local chromatin motion is maintained at a similar level throughout interphase under physiological conditions.

**Figure 10.**
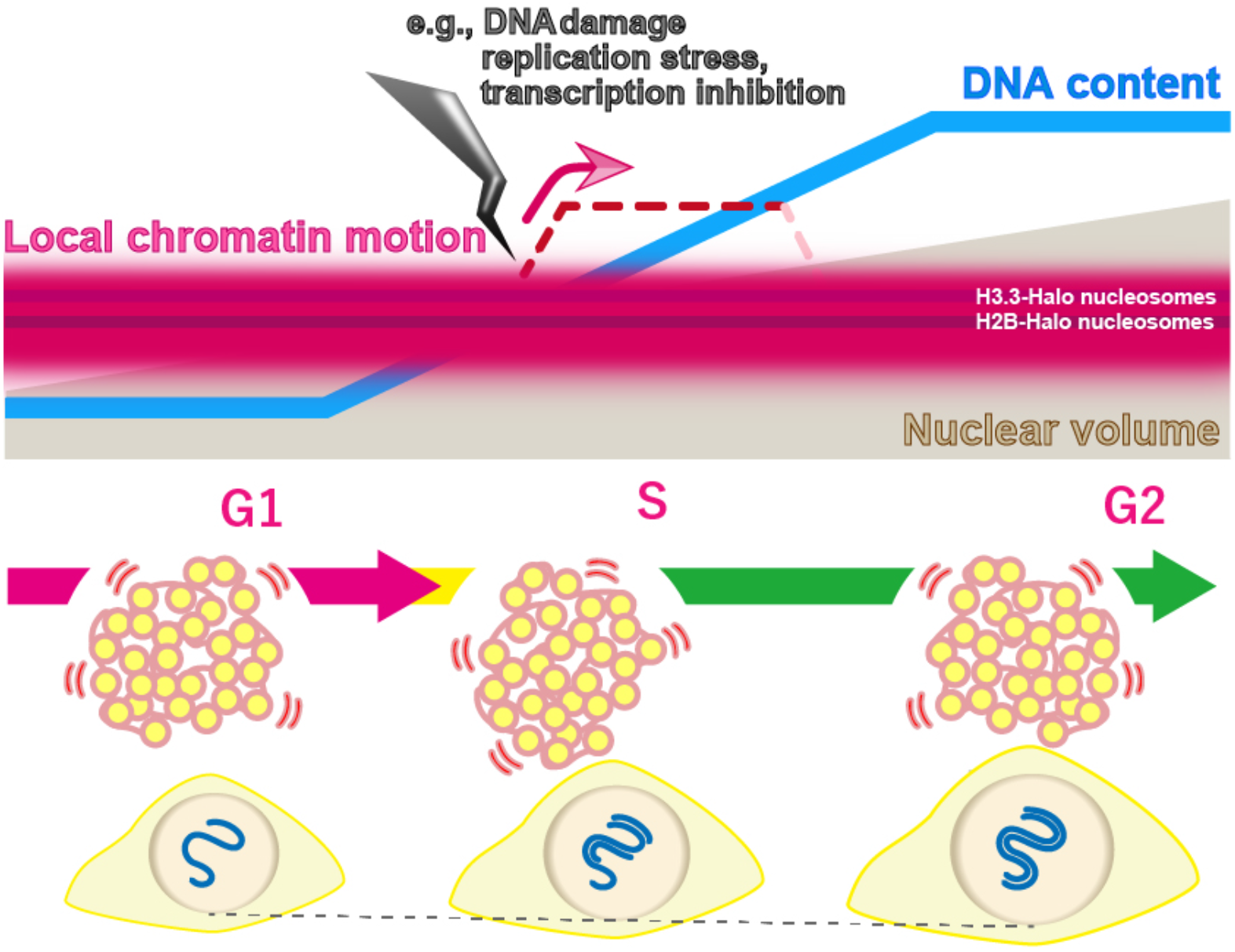
Model for local chromatin motion during interphase. Schematic summary of the present study. Local motion of nucleosomes (H2B-Halo nucleosomes) remains nearly constant from G1 through G2, although DNA content and nuclear volume increase during interphase. Local chromatin motion spans a range from more mobile euchromatin (H3.3-Halo nucleosomes) to less mobile heterochromatin, and this motion hierarchy is maintained throughout interphase. Local chromatin motion can be transiently increased by perturbations, such as DNA damage, replication stress, or transcription inhibition. This near-constant but tunable local motion may provide similar physical conditions for housekeeping genome functions during interphase.

This main conclusion is also consistent with our recent Repli-Histo labeling study [14, 27]. In the study, four known chromatin classes (Class IA, IB, II, and III) from euchromatin to heterochromatin were marked in living human cells and their local nucleosome motions were examined [14]. More euchromatic regions exhibited greater nucleosome motion, whereas more heterochromatic regions showed lower motion, and this relative motion hierarchy persisted throughout interphase (G1, S, and G2) [14]. Together with the present results including RL analysis (Fig. S3B) and H3.3-Halo-labeled nucleosomes, which are enriched in euchromatin (Figs. 4E and S4F), these findings suggest that interphase chromatin maintains a nearly constant population average of local motion while preserving heterogeneity in local nucleosome motion among chromatin classes, including euchromatin and heterochromatin (Fig. 10).

At first glance, our observations may appear inconsistent with previous reports showing that chromatin is more constrained in S and G2 than in G1 when analyzed on minute and micrometer or longer spatiotemporal scales [20–22]. However, this apparent difference can be understood from the scale-dependent viscoelastic nature of chromatin [5–8]. On larger spatial scales and longer time scales, chromatin behaves more solid-like, and each chromosome is quite stably occupied in its territory without excess intermingling and subsequent chromosome breaks, which contributes to maintaining genome integrity [7, 9]. In contrast, on subsecond and 100 to 300 nm scales, chromatin is locally more flexible and liquid-like [7]. Our results suggest that, on this local scale, nucleosome motion is maintained at a similar level from G1 through S and G2. This near-constant local nucleosome motion may help maintain comparable dynamic accessibility (penetrability) of chromatin throughout interphase [13, 14], thereby providing a stable physical environment in which housekeeping functions such as RNA transcription, DNA replication, and DNA repair can proceed [18]. On the other hand, larger-scale chromatin organization, such as A/B compartment organization, may change with interphase progression. Indeed, previous Hi-C and related chromosome conformation studies suggest that A/B compartment organization is re-established in early G1 and can continue to change during interphase, particularly in association with S-phase progression [70–72]. The considerable differences in response to transcription inhibition (Figs. 5B and 9) among G1, G1-S, and S-G2 phases may reflect such compartmental changes.

Our perturbation experiments further showed that local nucleosome motion can be modulated. Transcription inhibition by THZ1 increased nucleosome motion in all interphase phases examined, suggesting that RNA Pol II-dependent transcriptional activity normally contributes to constraining local chromatin motion, consistent with previous studies [52, 58–63] [64]. In addition, cellular perturbations, including replication stress induced by HU and DNA damage induced by cisplatin, also increased average nucleosome motion. Although the underlying mechanisms may differ among these perturbations, and remain unclear, all treatments shifted chromatin toward a more mobile state. Elevated chromatin motion upon replication stress and DNA damage may help recruit protein factors, such as DNA repair factors. Our findings suggest that cells can transiently change chromatin motion to perform ad hoc tasks in response to signals from inside and outside the cell.

Finally, what drives local nucleosome motion in interphase chromatin? We consider that local nucleosome fluctuation is primarily driven by thermal fluctuations, because a compact polymer model with thermal fluctuations closely recapitulates nucleosome motion in living cells [18, 73]. The extent of this motion is then restricted by chromatin context and associated factors. Nucleosome motion becomes increasingly constrained from euchromatin to heterochromatin during interphase [14, 27] and is most constrained in mitotic chromosomes [74]. As discussed above, transcription machinery likely contributes to constraining nucleosome motion in euchromatin, whereas HP1 and the nuclear lamina may play similar roles in heterochromatin. In mitotic chromosomes, condensins are likely to provide the major constraint. In this view, ATP-dependent active processes are unlikely to be a major direct driving force for nucleosome fluctuations, although ATP depletion seemed to increase chromatin dynamics [48, 75–77]. Notably, previous ATP depletion experiments are very likely to induce secondary effects, including chromatin condensation associated with increased free Mg^2+^, which can reduce chromatin motion [4, 18, 48, 78]. Systematic and rapid knockdown analyses of ATP-dependent chromatin proteins, such as remodelers, in cells would provide new insight into this issue.

### Limitations of this study

MSD analysis of H2B-Halo nucleosomes in HeLa cells showed that the early G1 phase had slightly higher MSD values, reaching statistical significance in comparisons with G1-S and S-G2 but not with G1 (Fig. 3A). This difference was not reproduced for H3.3-Halo nucleosomes in HeLa cells (Fig. 4E) or H2B-Halo nucleosomes in HCT116 cells (Fig. 8C). These results indicate that the slight increase in nucleosome motion in early G1 is not a robust feature. Further investigation is needed.

In addition, population-averaged analyses of single-nucleosome imaging using H2B-Halo or H3.3-Halo may still mask small differences. A time-lapse study tracking single nucleosomes throughout the cell cycle in live cells would be ideal, although improved photostability of fluorescent dyes would be required.

## Conclusions

Our results support a model in which local chromatin motion is maintained near a steady level throughout interphase under normal conditions, while remaining responsive to cellular perturbations, including transcriptional inhibition, replication stress, and DNA damage. This physical framework would help genome DNA carry out housekeeping functions in living human cells under broadly similar local conditions throughout interphase.

## Methods

### Establishment of stable cell lines

HeLa cells were cultured at 37 °C in 5% CO₂ in DMEM (D5796-500ML; Sigma-Aldrich) supplemented with 10% FBS (F7524; Sigma-Aldrich). HCT116 cells were cultured at 37 °C in 5% CO₂ in McCoy’s 5A medium (SH30200.01; HyClone) supplemented with 10% FBS.

For the establishment of HeLa cells stably expressing Fucci2, HeLa/Fucci2 cells were obtained from RIKEN BRC (RCB2867). To stably express H2B-Halo in the HeLa cell line, the PiggyBac transposon system was used. The constructed plasmid pPB-CAG-IB-H2B-HaloTag (Addgene #247345) was co-transfected with pCMV-hyPBase (provided by the Sanger Institute under a materials transfer agreement) into the HeLa cells using the Effectene transfection reagent kit (301425; Qiagen). Transfected cells were selected with 3 µg/mL blasticidin S (029-18701; Wako). For the establishment of Fucci(SA)5 HCT116 cells expressing H2B-Halo, HCT116 cells with endogenously tagged mAID-BromoTag-ORC1, mAID-BromoTag-CDC6 and OsTIR1(F74G) were used as parental cells [35]. A Fucci(SA)5 plasmid containing a puromycin selection marker was constructed from the tFucci(SA)5 Bsr plasmid by replacing the blasticidin S resistance gene with a puromycin resistance gene (Addgene #153520; [34]). This plasmid was transfected together with pCMV-hyPBase. Subsequently, the stable cells were selected in the presence of 1 µg/mL of puromycin (P8833-25MG; Sigma-Aldrich) for one week. The PA-mCherry region of pPB-EF1α-H2B-PA-mCherry-PGKneo (Addgene #247340) was replaced with HaloTag to make pPB-EF1α-H2B-HaloTag-PGKneo. pPB-EF1α-H2B-HaloTag-PGKneo was co-transfected with pCMV-hyPBase into the HCT116 cells and stable cells were selected with 700 µg/mL G418. To stably express H3.3-Halo in the HeLa Fucci2 cell line, pPB-EF1α-H3.3-HaloTag [51] was co-transfected with pCMV-hyPBase using the Effectene transfection reagent kit. Transfected cells were selected with 10 µg/mL blasticidin S.

### Expression and localization of H2B-HaloTag or H3.3-HaloTag

To examine the expression level of H2B-HaloTag, transfected cells were lysed in Laemmli sample buffer [79] supplemented with 10% 2-mercaptoethanol (133-1457; Fujifilm Wako) and incubated at 95 °C for 5 min to denature proteins. Cell lysates, equivalent to 1 × 10⁵ cells per lane, were subjected to 12.5% SDS–PAGE. For western blotting, the fractionated proteins in the gel were transferred to a PVDF membrane (IPVH00010; Millipore) using a semi-dry blotter (BE-320; BIO CRAFT). The membranes were blocked with 10% or 3% skim milk (190-12865; Fujifilm Wako) for detection of H2B or HaloTag, respectively. The membranes were then incubated with primary antibodies: rabbit anti-H2B (1:10,000; ab1790, Abcam) or mouse anti-HaloTag (1:1,000; G9211, Promega), followed by the corresponding goat horseradish peroxidase (HRP)-conjugated secondary antibodies: anti-rabbit IgG (1:5000; 170-6515, Bio-Rad) or anti-mouse IgG (1:5000; 170-6516, Bio-Rad). Bands were detected by chemiluminescence reactions (WBKLS0100; Millipore), and images were acquired with the EZ-Capture MG (AE-9300H-CSP; ATTO). To examine the expression level of H3.3-HaloTag, rabbit anti-H3 (1:100,000; ab1791, Abcam) and mouse anti-HaloTag (1:1,000; G9211, Promega) antibodies were used.

To examine the cellular localization of H2B-HaloTag or H3.3-HaloTag, cells grown on poly-L-lysine–coated coverslips (P1524-500MG; Sigma-Aldrich; C018001; Matsunami) were treated with 10 nM JF646 ligand (GA1120, Promega) overnight at 37 °C in 5% CO₂. The cells were fixed with 1.85% formaldehyde (064-00406; Wako) at room temperature for 15 min, permeabilized with 0.5% Triton X-100 (T-9284; Sigma-Aldrich) for 5 min, and stained with 0.5 μg/mL DAPI (10236276001; Roche) for 5 min. Samples were mounted with PPDI (20 mM HEPES [pH 7.4], 1 mM MgCl₂, 100 mM KCl, 78% glycerol, and 1 mg/mL para-phenylenediamine [695106-1G; Sigma-Aldrich]) [80]. Z-stack images (0.2 µm steps, 30 sections) were acquired using a DeltaVision Ultra microscope (GE Healthcare) equipped with an Olympus PlanApoN 60× objective (NA 1.42), an sCMOS camera, an InsightSSI light source (∼50 mW), and a four-color standard filter set. Generally, H2B-Halo shows genome-wide localization due to the frequent replacement of histone H2B within a few hours [40]. Expressed H3.3-Halo was localized to DAPI-poor regions in nuclei.

### Validation of Fucci2 or Fucci(SA)5 function

To confirm whether Fucci2 or Fucci(SA)5 in our cell lines function properly, cells were cultured on poly-L-lysine–coated coverslips at 37 °C in 5% CO₂. The cells were fixed with 1.85% formaldehyde at room temperature for 15 min, permeabilized with 0.5% Triton X-100 for 5 min, and stained with 0.5 μg/mL DAPI for 5 min. Samples were mounted with PPDI. Z-stack images (0.2 µm steps, 30 sections) were acquired using a DeltaVision Ultra microscope equipped with an Olympus PlanApoN 40× or 60× objective (NA 1.40 or 1.42, respectively), an sCMOS camera, an InsightSSI light source (∼50 mW), and a four-color standard filter set.

### Single-nucleosome imaging

Established cell lines were cultured on poly-L-lysine–coated glass-based dishes (3970-035; Iwaki). H2B-Halo or H3.3-Halo incorporated into nucleosomes was fluorescently labeled with 100 pM (HeLa) or 200 pM (HCT116) HaloTag JF646 ligand for 20 min at 37 °C in 5% CO₂, washed three times with 1× HBSS (H1387; Sigma-Aldrich), and then incubated in the following media overnight before single-nucleosome imaging. HeLa cells were observed in DMEM (21063-029; Thermo Fisher Scientific), and HCT116 cells in McCoy’s 5A (1-18F23-1; BioConcept). These media were phenol red–free and supplemented with 10% FBS.

A live-cell chamber (INU-TIZ-F1; Tokai Hit) and digital gas mixer (GM-8000; Tokai Hit) were used to maintain cell culture conditions (37 °C, 5% CO₂, and humidity) during microscopy. Single nucleosomes were observed using an inverted Nikon Eclipse Ti2 microscope equipped with a 100-mW Sapphire 646-nm laser (Coherent) and an sCMOS ORCA-Flash 4.0 or ORCA-Fusion BT camera (Hamamatsu Photonics). Live cells labeled with JF646 were excited with the 646-nm laser through an objective lens (100× PlanApo TIRF, NA 1.49; Nikon) and detected at 676–786 nm. An oblique illumination system with a TIRF unit (Nikon) was used to excite fluorescent nucleosome molecules within a thin area of the cell nucleus and reduce background noise (Fig. 2B). Fucci probes were imaged with light-emitting diode (LED) epi-illumination (X-Cite XYLIS, Excelitas Technologies) and detected at 503 to 547 nm and 582 to 636 nm. Sequential image frames were acquired using NIS-Elements (Nikon) at a frame rate of 50 ms under continuous illumination. Note that freely diffusing, non-nucleosomal histones cannot be tracked at this frame rate.

### Single-nucleosome tracking analysis

To study nucleosome motion within chromatin domains accurately, we mainly focused on the 0–0.5 s time window, which corresponds to the spatial range of typical chromatin domain sizes (up to ∼300 nm). At longer time scales, other factors, such as higher-order structures (e.g., compartments, territories) and nuclear movements, become more influential (for details, see [18, 81]).

Image processing, single-nucleosome tracking, and single-nucleosome movement analysis were performed as previously described [18, 48, 52, 82]. Briefly, background noise was subtracted using the rolling-ball background subtraction (radius, 50 pixels) in ImageJ. Nuclear regions in the images were manually extracted. Following this step, the centroid of each fluorescent dot in each image was determined by 2D Gaussian fitting and its trajectory was tracked with u-track, a MATLAB package [45].

To assess positional accuracy, we calculated the standard deviation of the two-dimensional movement of immobilized nucleosomes per 50 ms in FA-fixed HeLa and HCT116 cells (N = 10 nucleosomes). The localization accuracy was defined as the mean of SDx and SDy. Single-step photobleaching profiles confirmed that individual dots represented single nucleosomes.

For single-nucleosome imaging/tracking, we calculated displacement and mean squared displacement (MSD) of nucleosomes, because MSD captures the relevant constrained nucleosome dynamics, for the following reasons: the vast majority of the labeled histones are incorporated into nucleosomes constrained along a very long polymer rather than freely diffusing, and state mixing is minimal. In this respect, single-nucleosome imaging is very different from single-molecule tracking (SMT) of transcription factors (TFs). MSD in SMT of TFs can be error-prone because multiple diffusion states (free 3D diffusion, 1D sliding, specific binding) interconvert, and displacement-based analyses are often preferable (e.g., [83]).

For single-nucleosome movement analysis, the displacement and MSD of the fluorescent dots were calculated based on their trajectory using a Python script. In our tracking, trajectories of single nucleosome dots on the XY plane 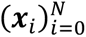 were acquired, where ***x****_i_* indicates XY-coordinates at the time point *i*. Then, we calculated the MSD for the lag time Δ by:

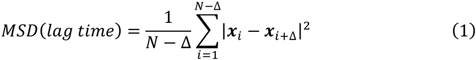

The originally calculated MSD was in 2D. To obtain the 3D value, the 2D value was multiplied by 1.5 (4Dt to 6Dt). The calculated MSDs were fitted to a subdiffusive model *MSD*(*t*) = 6*D* · *t^α^*, where α is an anomalous diffusion exponent (0 < α < 1). Statistical analyses of the obtained single-nucleosome MSD values for each histone protein were performed using Python. For Spot-On analysis [83], single-nucleosome trajectory data acquired at 10 ms/frame were analyzed. A three-state model was used for model fitting.

### Chemical treatment in single-nucleosome imaging

For chemical fixation, cells grown on poly-L-lysine coated glass-based dishes were incubated in 2% formaldehyde (diluted pure 16% formaldehyde) in 1 × HBSS at room temperature for 15 min and washed with 1 × PBS. For transcription inhibition, cells were treated with 1 µM THZ1 (CS-3168; Funakoshi) for 2 h. For induction of replication stress or DNA damage, cells were incubated in medium supplemented with 1 mM hydroxyurea (H8627; Sigma) or 4 µM cisplatin (033-20091; Wako) for 24 h. HU and cisplatin took longer to produce detectable effects on nucleosome motion. Control cells were treated with 0.1% DMSO (solvent used to dissolve THZ1), 0.2% Milli-Q water (solvent used to dissolve hydroxyurea) or 0.08% 1×PBS (used to dissolve cisplatin). After these treatments, cells were live-cell imaged or chemically fixed.

### Nuclear volume measurement

Nuclear volumes were measured as described previously [18]. To measure nuclear volume, the cells were grown on poly-L-lysine–coated coverslips. Subsequent processes were performed at room temperature. The cells were fixed with 1.85% FA for 15 min, permeabilized with 0.5% Triton X-100 for 15 min, and stained with DAPI (0.5 µg/mL) for 5 min, followed by PPDI mounting. Z-stack images (every 0.4 µm in the z direction, 55 sections in total) of the cells were obtained using a FLUOVIEW FV3000 confocal laser scanning microscope (OLYMPUS) equipped with an Olympus UPLANSAPO 60×W objective [numerical aperture (NA), 1.20] at room temperature. Obtained Z-stack images were loaded to Imaris 10.0.1 (Bitplane AG, Zurich, Switzerland) and converted to Imaris 3D image files (.ims). To calculate the nuclear volume, the tool “Surface” was used. A threshold value was automatically (HeLa cells) and manually (HCT116 cells) determined to include all DAPI signals. Note that only well-isolated nuclei were recorded and analyzed.

### Indirect immunofluorescence

Immunostaining was performed as described previously [13], and all processes were performed at room temperature. Cells were fixed in 1.85% FA in PBS for 15 min and then treated with 50 mM glycine in HMK [20 mM Hepes (pH 7.5) with 1 mM MgCl_2_ and 100 mM KCl] for 5 min and permeabilized with 0.5% Triton X-100 in HMK for 5 min. After washing twice with HMK for 5 min, the cells were incubated with 10% normal goat serum (NGS; 143-06561, Wako) in HMK for 30 min. The cells were incubated with the diluted primary antibody, rabbit anti-gamma H2A.X (1:1000; ab2893, Abcam) or mouse anti-RNAP II Ser5P (1:1000; provided by Kimura Lab, Science Tokyo, Japan), in 1% NGS in HMK for 1 h. After being washed with HMK four times, the cells were incubated with the diluted secondary antibody, goat anti-rabbit IgG Alexa Fluor 647 (1:1000; A21245, Invitrogen) or goat anti–mouse IgG Alexa Fluor 488 (1:500; A11029, Invitrogen), in 1% NGS in HMK for 1 h followed by a wash with HMK four times. For DNA staining in fixed cells, DAPI (0.5 μg/mL) was added to the cells for 5 min followed by washing with HMK. The stained cells were embedded in PPDI.

### Fluorescence microscopy on fixed samples

Image stacks of fixed cells were acquired by using Delta Vision Ultra (GE Healthcare) with an Olympus UPLXAPO 100× (NA 1.45) or UPLXAPO 60× (NA 1.42) objective. Optical sections were acquired at 0.2 μm intervals. For interphase nuclei, the best-focused section near the middle of each nucleus was extracted after chromatic aberration correction. Image analysis was performed using Fiji/ImageJ.

To quantify intranuclear signal intensities, DAPI-stained nuclear regions were segmented using Huang’s fuzzy thresholding method in Fiji/ImageJ, and the mean pixel intensities (a.u.) within the nuclear regions were measured for DAPI and all other channels.

### Bayesian-based Richardson-Lucy (RL) algorithm

The RL algorithm is the iterative algorithm of Richardson [84] and Lucy [85] (RL) to derive smooth distributions from the noisy data used in image processing [86, 87]. With sufficient observation, the algorithm converts raw, noisy trajectories into a smooth distribution of MSD, providing a denoised, population-level view of heterogeneous nucleosome mobility.

For the convenience of the reader, we briefly explain the mathematical principles of the RL algorithm. A more detailed description has been reported previously [11].

Here, the average MSD of nucleosomes,*M̄* = ⟨*M_i_*⟩, where 〈⋯ 〉 is the average taken over *i* and along the observed trajectories.

The distribution of MSD is captured by

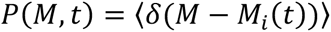

Due to the short lifetime of observed fluorescence of single nucleosomes, the individual nucleosome MSD data are insufficient to provide for a clear *P*(*M*, *t*). However, this problem is overcome by using the RL algorithm. From the observed data, we first calculate the self-part of the van Hove correlation function (vHC),

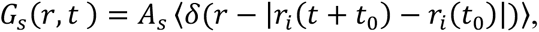

where *r_i_* is the projected coordinate of the *i*th nucleosome on the 2-dimensional imaging plane and *A_s_*is a constant to normalize *G_s_* as ∫ *d*^2^ *rG_s_*(*r*, *t*) = 1. The calculated vHC is shown at *t* =0.5 s. *G_s_* is expanded in Gaussian bases 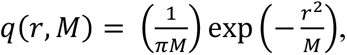 as

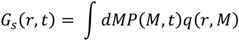

Given a noisy estimate of *G_s_*(*r*, *t*)*, P*(*M*, *t*) is extracted as coefficients of expansion using the RL iterative scheme. For this, *P*(*M*, *t*) was calculated in an iterative way with the RL algorithm: starting from the initial distribution, 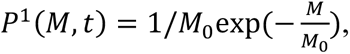, *P^n^*^+1^(*M*, *t*) at the (n+1)-th iteration was obtained by 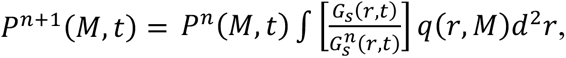 This equation was iterated under the constraints *P^n^*^+1^(*M*, *t*) > 0 and ∫ *P^n^*^+1^ (*M*, *t*)*dM* = 1.

The obtained distribution curves were fitted with multiple Gaussian functions, and the peak position and area of each Gaussian component were calculated. Subpopulation boundaries were estimated from the RL-curve: using the local minima for the “Super slow”–“Slow” and “Slow”–“Fast” transitions, and the midpoint between peaks for the “Fast”–“Super fast” transition. For MSD analysis, trajectories within each cell were categorized based on their MSD at Δt = 0.5 s, and mean squared displacement (MSD) was calculated separately for each subpopulation.

### Genomic analysis of H2B-HaloTag nucleosomes in HeLa and HCT116 cells and H3.3-HaloTag nucleosomes in HeLa cells

The pull-down sequencing data of H2B-Halo nucleosomes and H3.3-Halo nucleosomes are publicly available in the DDBJ BioProject database under accession numbers PRJDB17378 (HeLa H2B-HaloTag; SRA runs DRR527334–DRR527339; https://ddbj.nig.ac.jp/search/entry/bioproject/PRJDB17378/)[88], PRJDB17380 (HeLa H3.3-HaloTag; SRA runs DRR527328–DRR527333; https://ddbj.nig.ac.jp/search/entry/bioproject/PRJDB17380/)[51], and PRJDB39674 (HCT116 H2B-HaloTag; SRA runs DRR801689–DRR801696; https://ddbj.nig.ac.jp/search/entry/bioproject/PRJDB39674/)[51]. Hi-C domains from HeLa [89] were obtained from GEO (GSE63525). Compartment annotations were from SNIPER predictions [90]. ChIP-seq peaks for histone modifications were downloaded from ENCODE [91, 92]: H3K4me2 (ENCFF108DAJ), H3K4me3 (ENCFF447CLK), H3K9ac (ENCFF723WDR), H3K27ac (ENCFF144EOJ), H3K79me2 (ENCFF916VLX), H3K36me3 (ENCFF001SVY), H3K4me1 (ENCFF162RSB), and H3K27me3 (ENCFF252BLX).

### Use of AI tools

OpenAI ChatGPT was used to assist with English-language proofreading and minor grammatical corrections. All suggested changes were reviewed by the authors, who take full responsibility for the manuscript.

## Acknowledgments

We are grateful to Dr. K. M. Marshall for critical reading of this manuscript. We thank Mr. H. Ogawa for initial results, Dr. K. Minami for critical reading and technical assistance, and Dr. S. Ide for technical assistance, Dr. K. Higashi, Dr. A. Toyoda, and Dr. K. Kurokawa for genomics analysis. We thank Dr. A. Miyawaki for providing Fucci2 cells and Fucci(SA)5 plasmids and Dr. H. Kimura for providing RNA Pol II antibodies. We also thank Dr. M. Sasai and Dr. S. S. Ashwin for development of RL analysis. Finally, we thank all members of the Maeshima laboratory for valuable discussions and continuous support.

## Author contributions

Y.N., S.I., and K.M. designed the research. Y.N. and S.I. performed the experiments including cell generation, imaging, and analyses. M.A.S. established RL analysis. S.T., K.N., and K.S. performed some biochemical experiments. Y.H. and M.T.K. contributed to cell generation. Y.N., S.I., M.A.S. and K.M. wrote the manuscript with help of K.S. with input from all authors.

## Ethics declaration

Not applicable.

## Data and materials availability

The scripts for RL-algorithm classification and single-nucleosome tracking analysis are available at https://doi.org/10.5281/zenodo.16959115. All data needed to evaluate the conclusions in the paper are present in the paper and/or the Supplementary Materials. Requests for the reagents, data, and scripts should be submitted to K.M.

---

**Figure S1.**
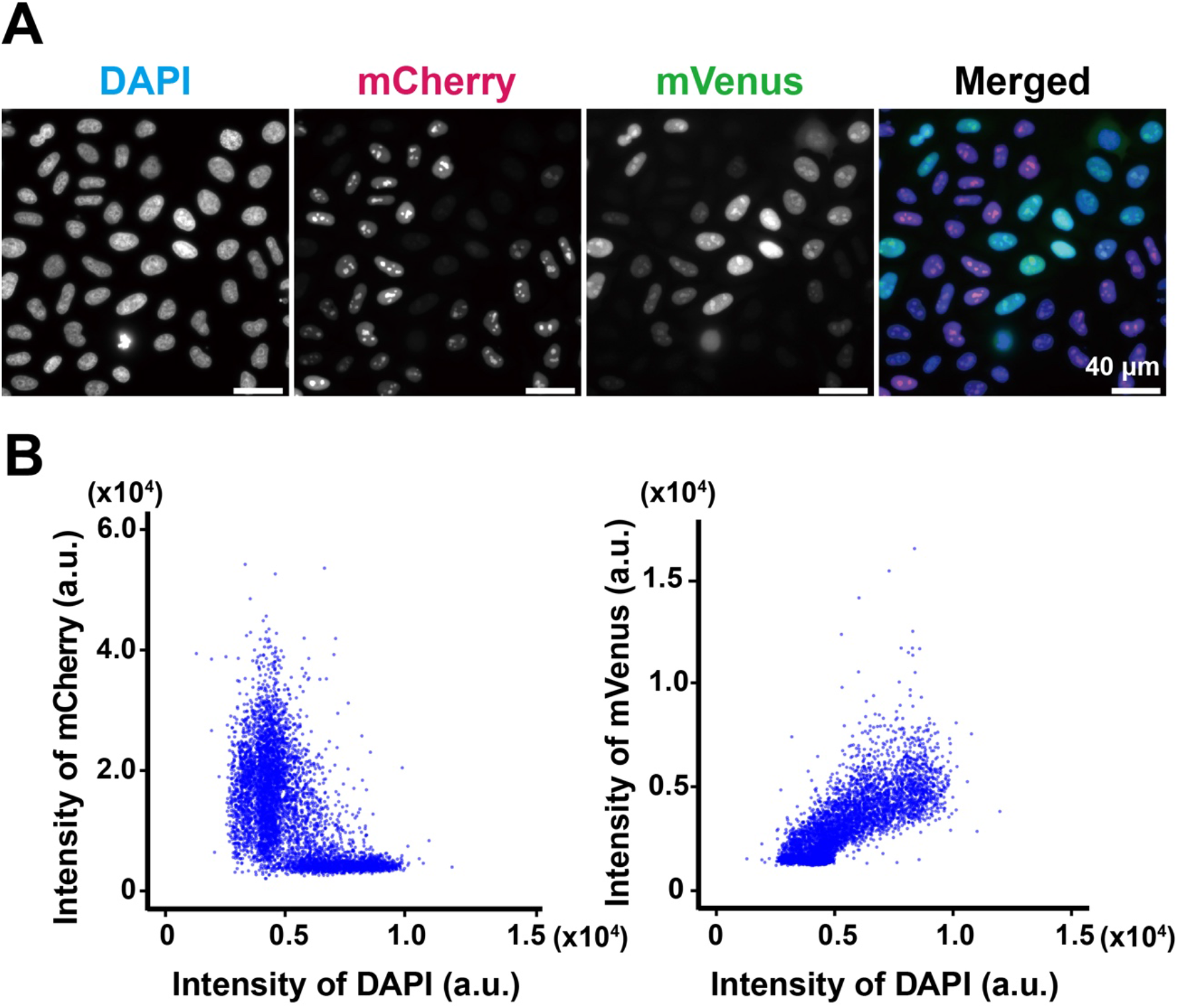
Fluorescence properties of Fucci2 probes in formaldehyde-fixed HeLa cells. **(A)** Representative images of DAPI-stained DNA, mCherry-hCdt1, and mVenus-hGeminin in formaldehyde-fixed HeLa Fucci2 cells. A merged image is shown on the right. Scale bars, 40 μm. **(B)** Scatter plots of mCherry-hCdt1 fluorescence intensity versus DAPI fluorescence intensity (left) and mVenus-hGeminin fluorescence intensity versus DAPI fluorescence intensity (right) in formaldehyde-fixed HeLa cells. mCherry-hCdt1 fluorescence was higher in cells with lower DAPI intensity, whereas mVenus-hGeminin fluorescence increased with increasing DAPI intensity, consistent with cell-cycle progression from G1 to S and G2 (n = 5547 cells).

**Figure S2.**
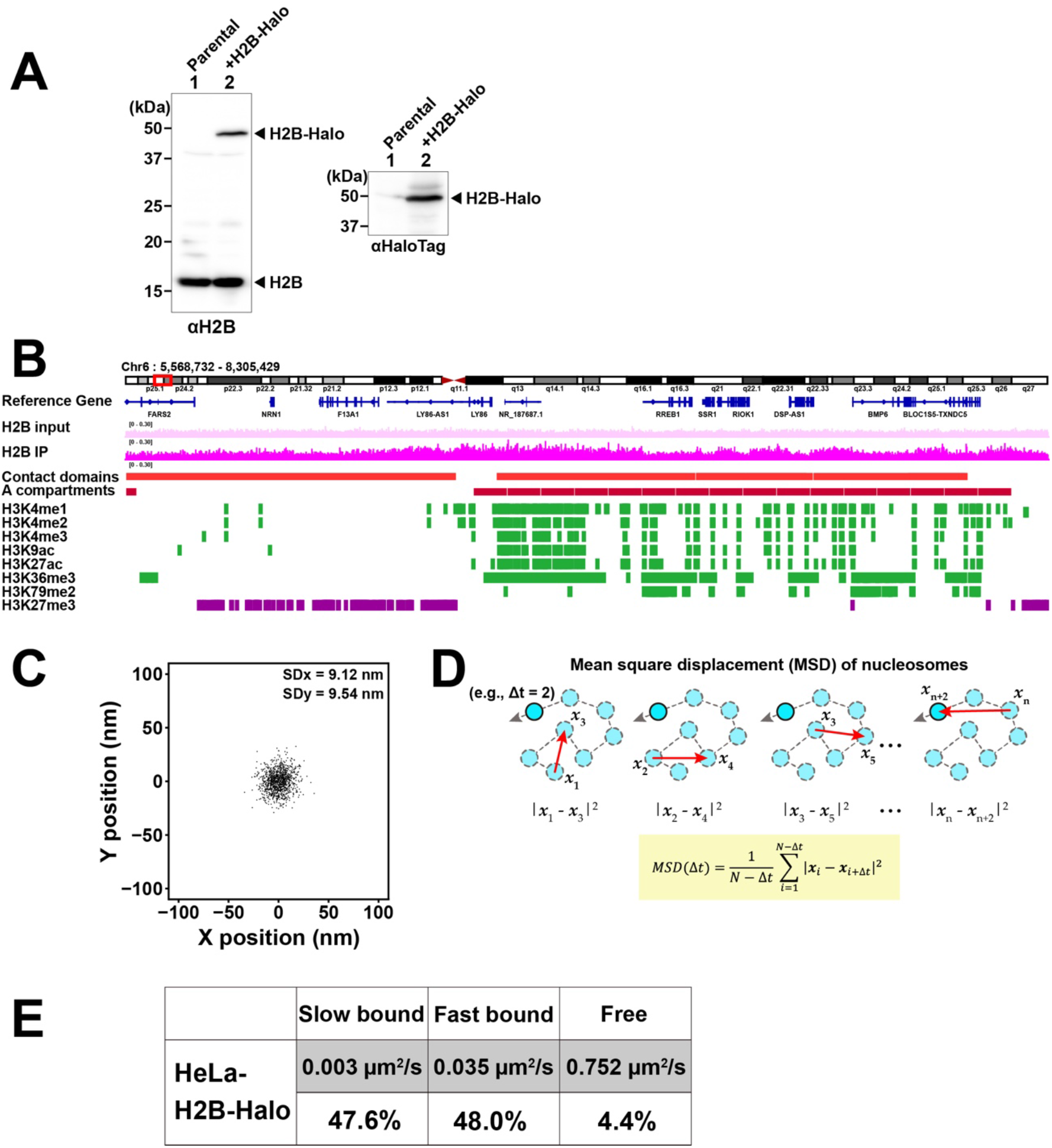
Characterization of H2B-HaloTag for single-nucleosome imaging. **(A)** Immunoblot of H2B-HaloTag-expressing cells and control cells probed with antibodies against HaloTag and histone H2B. **(B)** Genome browser view showing H2B-HaloTag incorporation across chromosome 6. H2B input and H2B immunoprecipitation signals are shown together with contact domains, A compartment, and representative histone modification profiles, indicating genome-wide incorporation of H2B-HaloTag into chromatin. Genomic data were reproduced from [88]. **(C)** Two-dimensional position distribution of immobilized fluorescent signals used to estimate localization precision (n = 10 nucleosomes). SDx and SDy are indicated. **(D)** Schematic illustration of mean square displacement (MSD) analysis of single-nucleosome trajectories. MSD was calculated from the squared displacements between positions separated by a given time interval, Δt. **(E)** Classification of H2B-HaloTag motion states by Spot-On [83]. Three motional states, slow bound, fast bound, and free, were identified, and their apparent diffusion coefficients and fractions are shown. Nucleosome trajectories used per cell: 1480 ∼ 5524; n = 31 cells.

**Figure S3.**
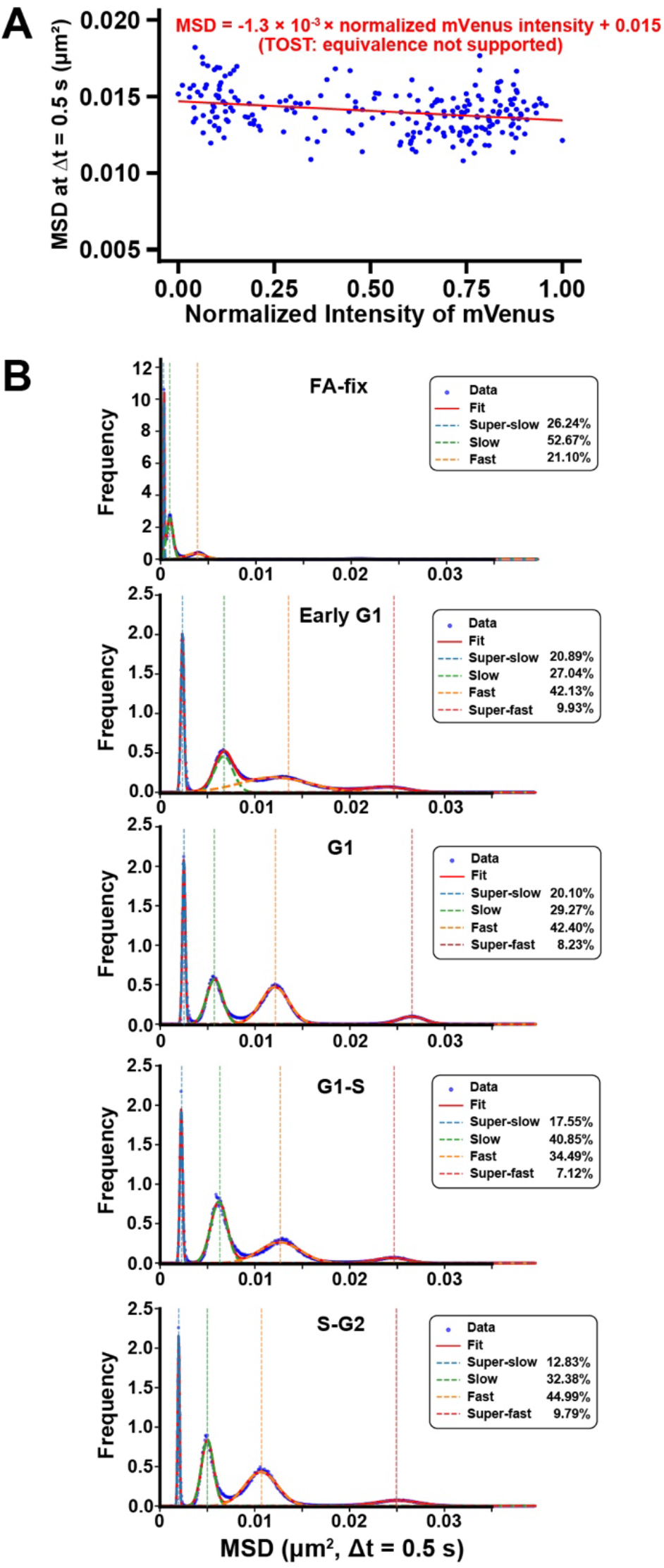
Additional analyses of local nucleosome motion throughout interphase. **(A)** Scatter plot of MSD at Δt = 0.5 s versus normalized mVenus fluorescence intensity for all cells, including early G1 cells. A weak negative trend was observed across the full range of mVenus intensity, reflecting the slightly elevated MSD values in early G1 cells. n = 226 cells. **(B)** Richardson-Lucy (RL) deconvolution analysis of MSD distributions at Δt = 0.5 s in early G1 (n = 35 cells), G1 (n = 28 cells), G1-S (n = 82 cells), and S-G2 (n = 78 cells) cells. Four motional components, designated super-slow, slow, fast, and super-fast, were identified in all cell-cycle phases. The overall distribution profiles were similar among G1, G1-S, and S-G2 cells, whereas early G1 cells showed a modest shift toward higher mobility. A control plot of formaldehyde-fixed cells is presented at the top.

**Figure S4.**
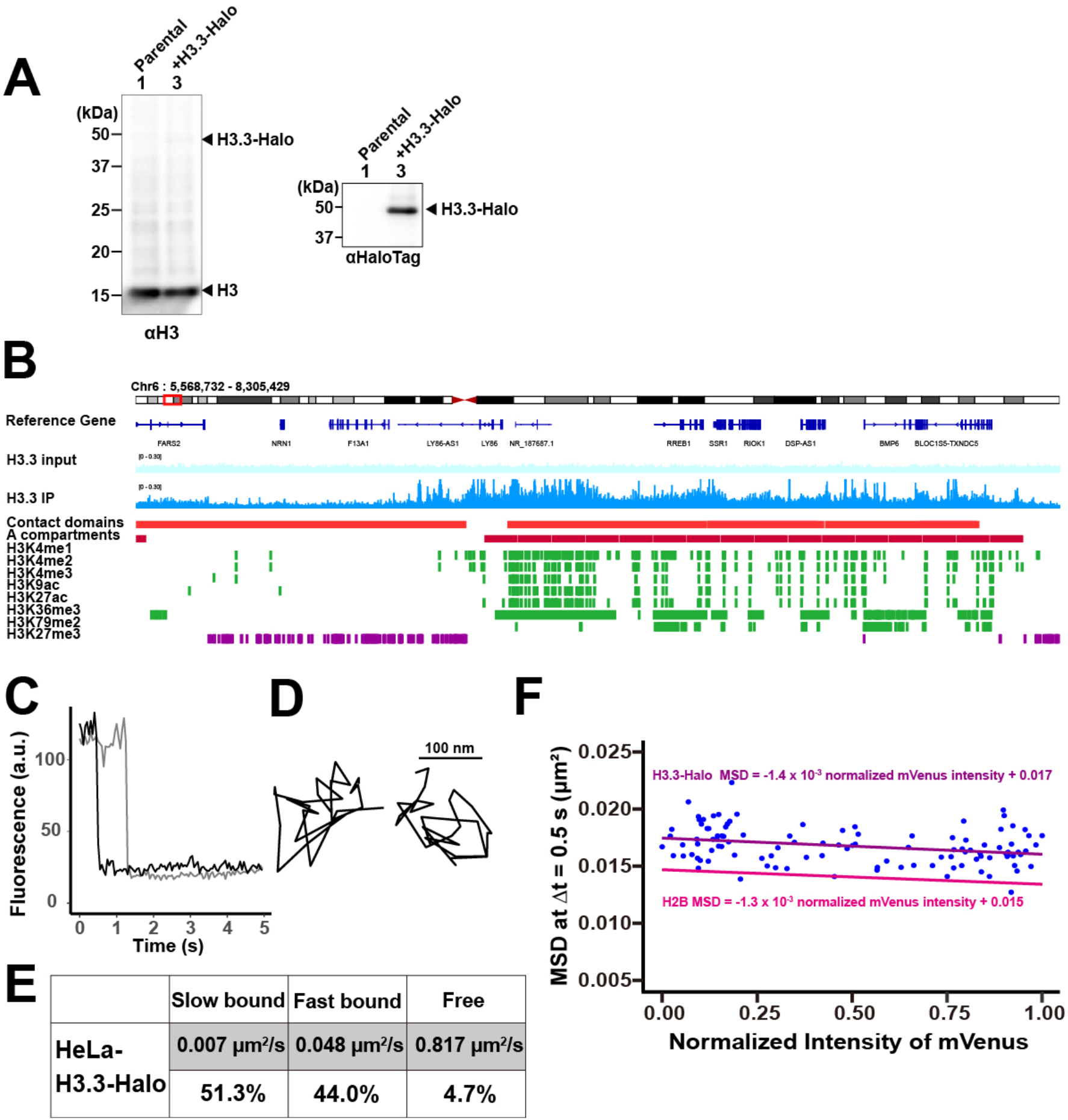
Characterization of H3.3-Halo for single-nucleosome imaging. **(A)** Immunoblot of parental HeLa Fucci2 cells and HeLa Fucci2 cells expressing H3.3-Halo, probed with antibodies against histone H3 and HaloTag. Arrowheads indicate H3.3-Halo and endogenous histone H3. **(B)** Genome browser view of H3.3-Halo incorporation across chromosome 6. H3.3 input and H3.3 immunoprecipitation signals are shown together with contact domains, A compartments, and representative active and repressive histone modification profiles. The genomic data were reproduced from [51]. **(C)** Representative fluorescence intensity traces of single H3.3-Halo-JF646 dots showing single-step photobleaching. a.u., arbitrary units. **(D)** Two representative trajectories of single H3.3-Halo nucleosomes in a living HeLa Fucci2 cell. Scale bar, 100 nm. **(E)** Spot-On analysis of H3.3-Halo molecules using a three-state model. The estimated diffusion coefficients and fractions of the slow-bound, fast-bound, and free populations are shown. Nucleosome trajectories used per cell: 1503 ∼ 5151; n = 31 cells. **(F)** Scatter plot of H3.3-Halo nucleosome MSD at Δt = 0.5 s versus normalized mVenus fluorescence intensity. Linear regression lines for H3.3-Halo and H2B-Halo nucleosomes are shown. The regression equations were MSD = −1.4 × 10⁻³ × normalized mVenus intensity + 0.017 for H3.3-Halo and MSD = −1.3 × 10⁻³ × normalized mVenus intensity + 0.015 for H2B-Halo. H3.3-Halo: n = 108 cells; H2B-Halo: n = 226 cells.

**Figure S5.**
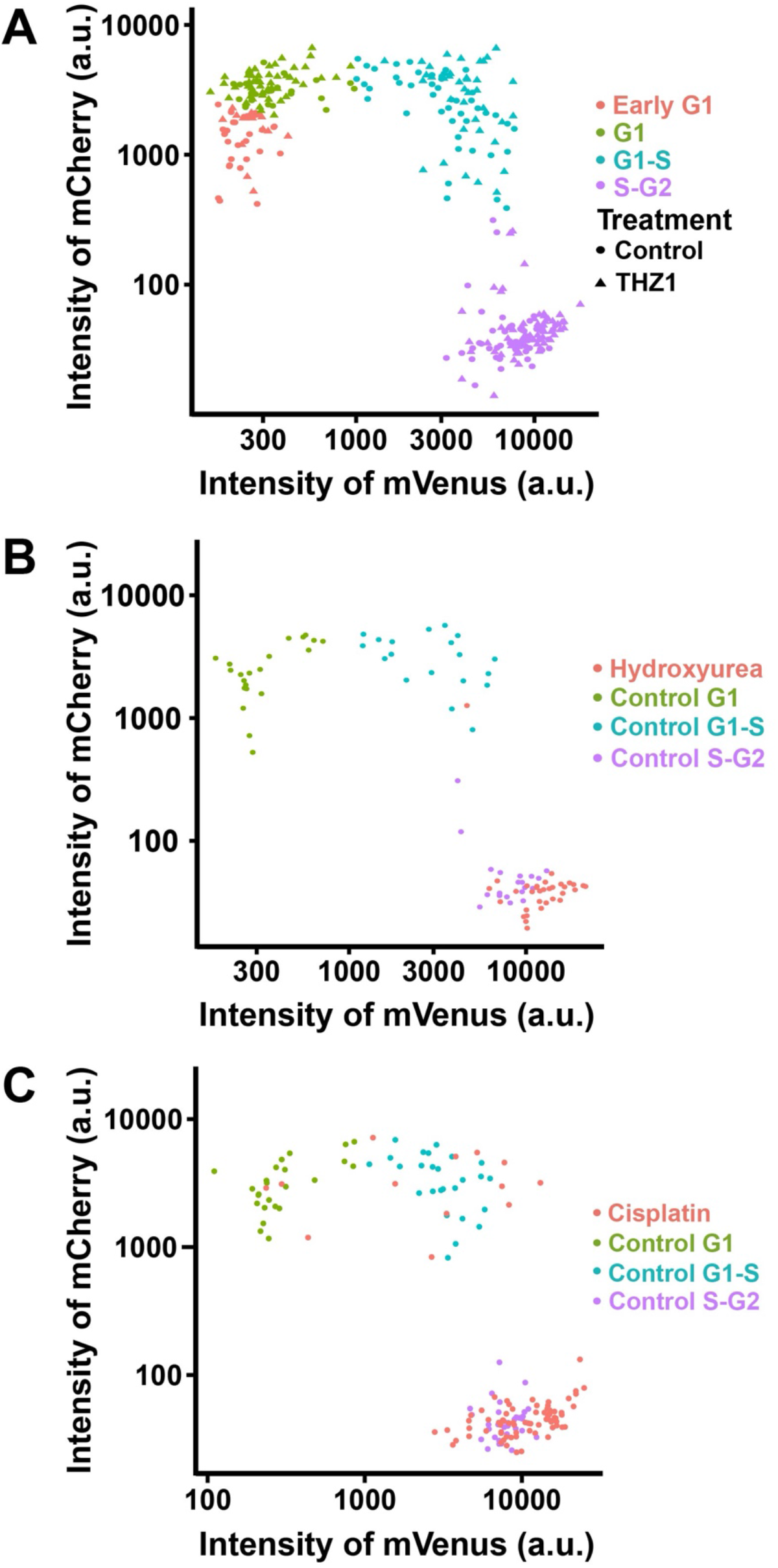
Cell-cycle distribution of drug-treated HeLa cells based on Fucci2 fluorescence intensities. **(A)** Scatter plots of mCherry-hCdt1 and mVenus-hGeminin fluorescence intensities in control and THZ1-treated (2 h) cells. Overlay of control and THZ1-treated cells. Cell-cycle populations are indicated by color, and treatments are distinguished by symbol shape. (Control: Early G1; n = 29 cells, G1; n = 24 cells, G1-S; n = 46 cells, S-G2; n = 53 cells. THZ1: Early G1; n = 15 cells, G1; n = 52 cells, G1-S; n = 39 cells, S-G2; n = 60 cells). **(B)** Scatter plot of hydroxyurea-treated (24 h) cells together with the corresponding control cell-cycle populations (G1, G1-S, and S-G2). Hydroxyurea-treated cells accumulated mainly in the S-phase-related populations. Control G1: n = 32 cells, Control G1-S: n = 27 cells, Control S-G2: n = 28 cells, Hydroxyurea: n = 59 cells. **(C)** Scatter plot of cisplatin-treated (24 h) cells together with the corresponding control cell-cycle populations. Cisplatin-treated cells accumulated predominantly in the S-G2 population. Control G1: n = 24 cells, Control G1-S: n = 26 cells, Control S-G2: n = 32 cells, Cisplatin: n = 83 cells (G1: n = 3 cells, G1-S: n = 10 cells, S-G2: n = 70 cells).

**Figure S6.**
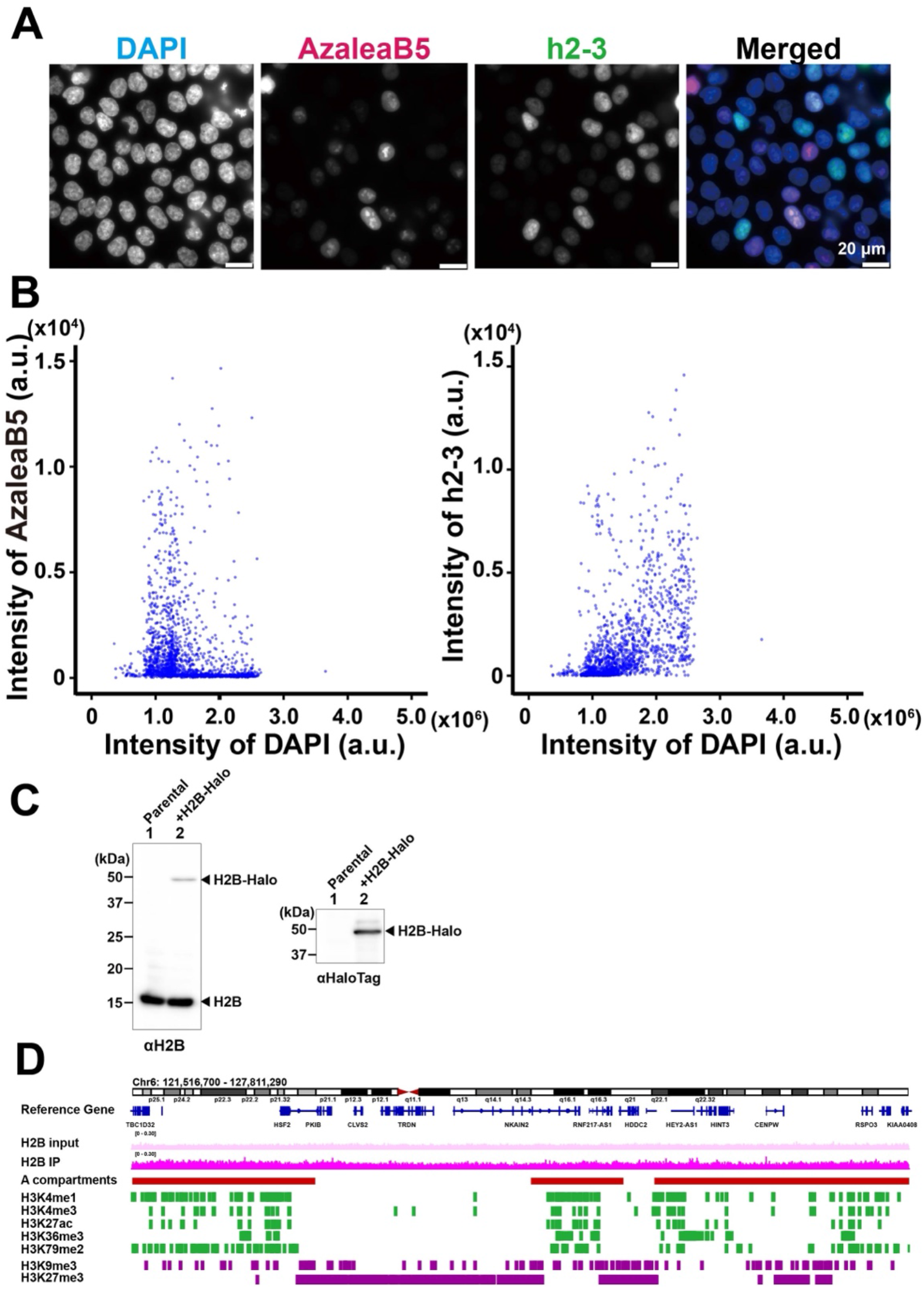
Characterization of Fucci(SA)5 probes and H2B-HaloTag expression in HCT116 cells. **(A)** Representative images of DAPI-stained DNA, AzaleaB5-hCdt1, and h2-3-hGeminin in formaldehyde-fixed HCT116 cells expressing Fucci(SA)5 probe. A merged image is shown on the right. Scale bars, 20 μm. **(B)** Scatter plots of AzaleaB5-hCdt1 fluorescence intensity versus DAPI fluorescence intensity (left) and h2-3-hGeminin fluorescence intensity versus DAPI fluorescence intensity (right) in formaldehyde-fixed HCT116 cells. n = 1735 cells. **(C)** Immunoblot of H2B-HaloTag-expressing HCT116 cells and control cells probed with antibodies against HaloTag and histone H2B. **(D)** Genome browser view showing H2B-HaloTag incorporation across chromosome 6. H2B input and H2B-Halo pull-down signals are shown together with contact domains, A compartment, and representative histone modification profiles, indicating genome-wide incorporation of H2B-HaloTag into chromatin. Genomic data were reproduced from [51].

**Figure S7.**
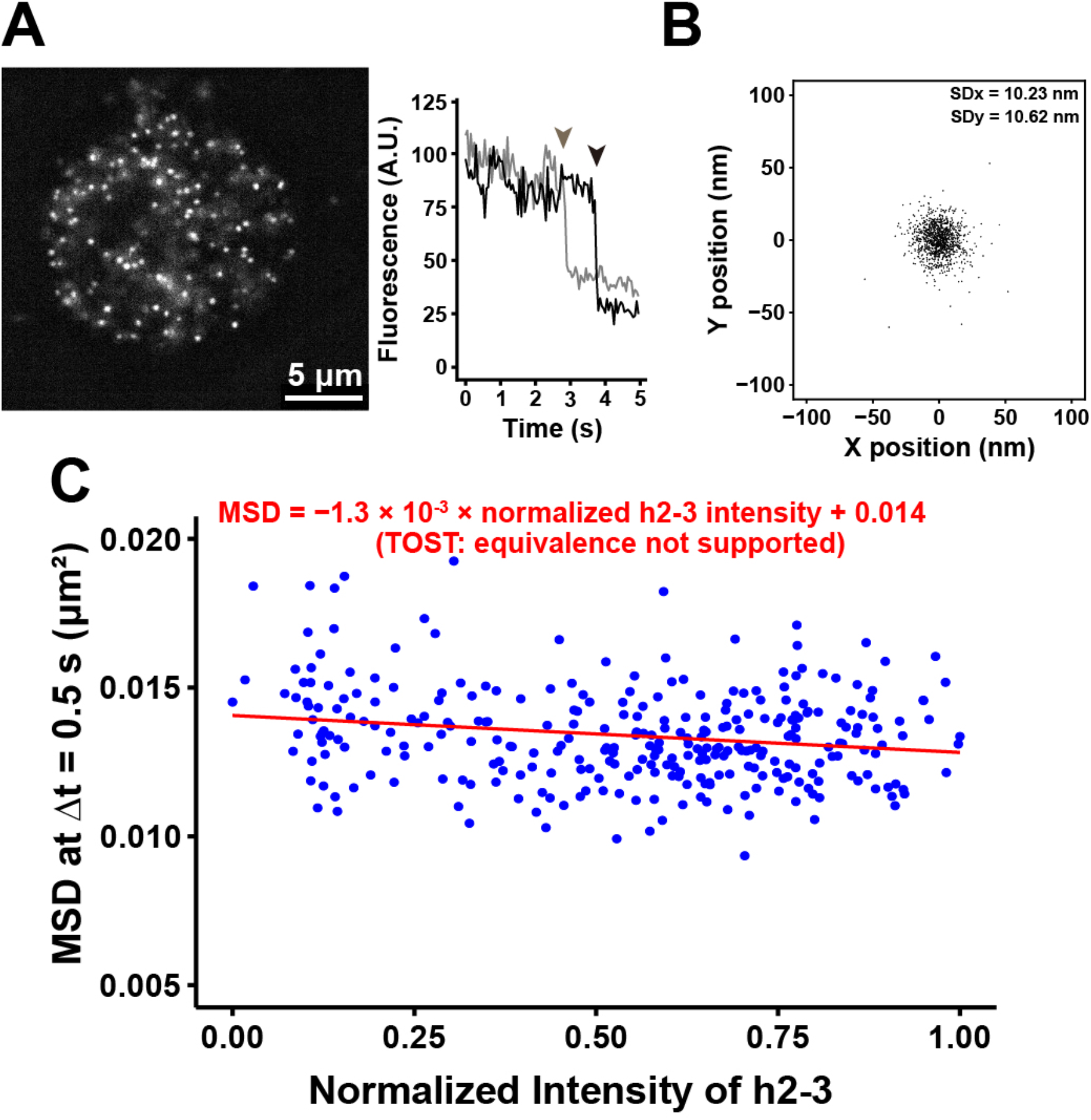
Additional analyses of HCT116 cells. **(A)** Representative image of single H2B-HaloTag-JF646 dots in the nucleus of a living HCT116 cell and representative fluorescence intensity traces showing single-step photobleaching (marked with arrow heads). **(B)** Two-dimensional position distribution of immobilized fluorescent signals used to estimate localization precision (n = 10 nucleosomes). SDx and SDy are indicated. **(C)** Scatter plot of MSD at Δt = 0.5 s versus normalized h2-3 fluorescence intensity for all HCT116 cells, including early G1 cells. A weak negative trend was observed across the full range. n = 302 cells.

**Figure S8.**
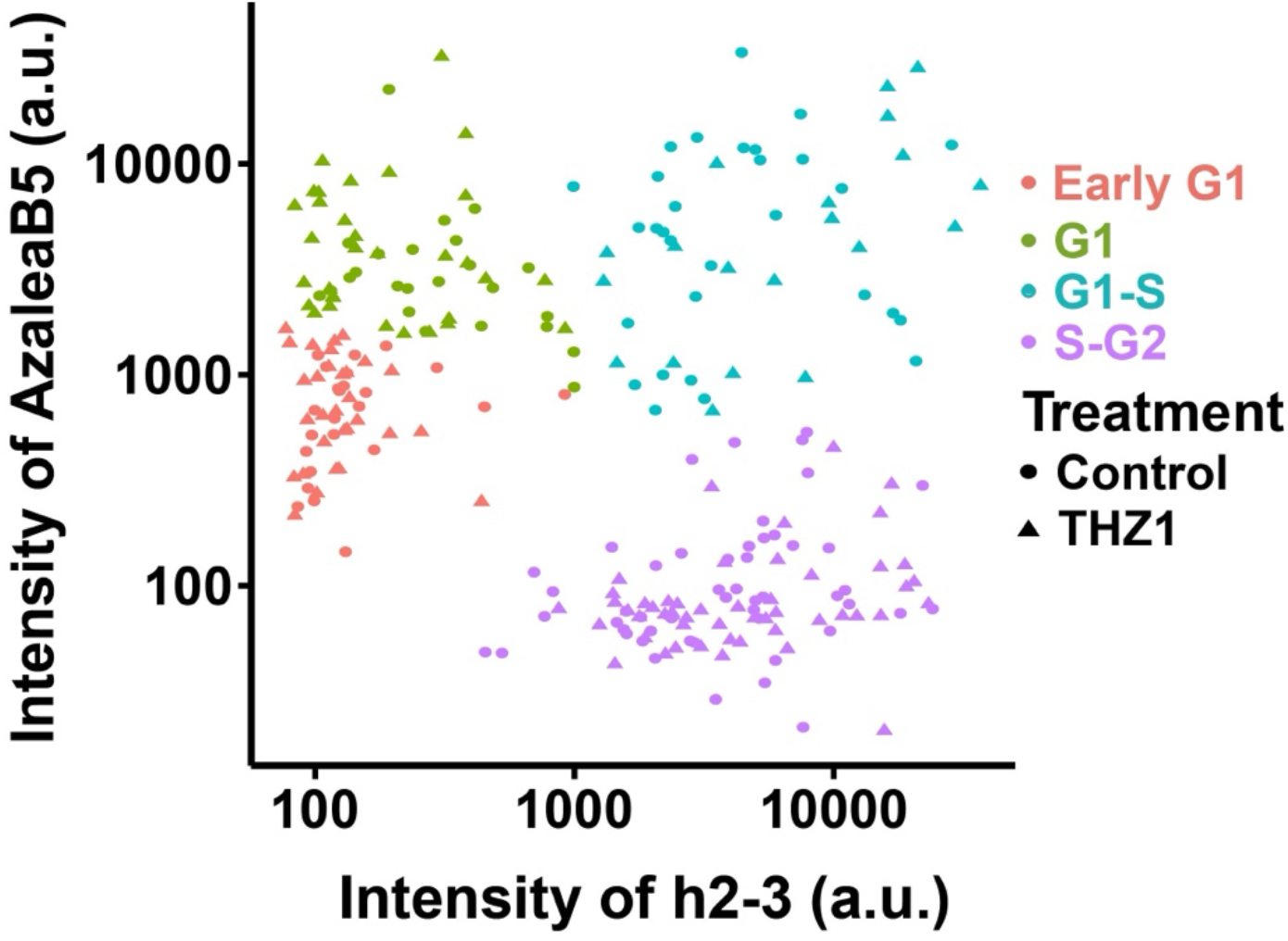
Classification of HCT116 cell-cycle populations based on Fucci(SA)5 fluorescence intensities. Scatter plot of AzaleaB5-hCdt1 and h2-3-hGeminin fluorescence intensities in HCT116 cells. Cells were classified into four cell-cycle populations, early G1, G1, G1-S, and S-G2, and treatments are distinguished by symbol shape, showing that THZ1 treatment (1 µM THZ1 for 2 h) did not markedly alter the overall Fucci(SA)5-based cell-cycle classification. (Control: Early G1; n = 24 cells, G1; n = 24 cells, G1-S; n = 30 cells, S-G2; n = 50 cells. THZ1: Early G1; n = 30 cells, G1; n = 32 cells, G1-S; n = 20 cells, S-G2; n = 51 cells).

## Movie legends

**Movie S1.**

Movie data (50 ms/frame) of single nucleosomes labeled with H2B-Halo-JF646 in a living HeLa Fucci2 cell recorded by sCMOS ORCA-Fusion BT camera (Hamamatsu Photonics). Note that clear and well-separated dots are visualized with a single-step photobleaching profile after background subtraction and contrast enhancement.

**Movie S2.**

Tracking data obtained by u-track (MATLAB package; [45]) were overlaid on Movie S1 and marked with circles.

**Movie S3.**

Movie data (50 ms/frame) of single nucleosomes labeled with H3.3-Halo-JF646 in a living HeLa Fucci2 cell recorded by sCMOS ORCA-Fusion BT camera (Hamamatsu Photonics). Note that clear and well-separated dots are visualized with a single-step photobleaching profile after background subtraction and contrast enhancement.

**Movie S4.**

Movie data (50 ms/frame) of single nucleosomes labeled with H2B-Halo-JF646 in a living HCT116 Fucci(SA)5 cell recorded by sCMOS ORCA-Fusion BT camera (Hamamatsu Photonics). Note that clear and well-separated dots are visualized with a single-step photobleaching profile after background subtraction and contrast enhancement.

**Movie S5.**

Tracking data obtained by u-track (MATLAB package; [45]) were overlaid on Movie S4 and marked with circles.

